# Primordial emergence of a nucleic acid binding protein via phase separation and statistical ornithine to arginine conversion

**DOI:** 10.1101/2020.01.18.911073

**Authors:** Liam M. Longo, Dragana Despotović, Orit Weil-Ktorza, Matthew J. Walker, Jagoda Jabłońska, Yael Fridmann-Sirkis, Gabriele Varani, Norman Metanis, Dan S. Tawfik

**Author notes:** These authors contributed equally to the work (Liam M. Longo, Dragana Despotović, Orit Weil-Ktorza).

## Abstract

*De novo* emergence, and emergence of the earliest proteins specifically, demands a transition from disordered polypeptides into structured proteins with well-defined functions. However, can peptides confer evolutionary relevant functions, let alone with minimal abiotic amino acid alphabets? How can such polypeptides evolve into mature proteins? Specifically, while nucleic acids binding is presumed a primordial function, it demands basic amino acids that do not readily form abiotically. To address these questions, we describe an experimentally-validated trajectory from a phase-separating polypeptide to a dsDNA-binding protein. The intermediates comprise sequence-duplicated, functional proteins made of only 10 amino acid types, with ornithine, which can form abiotically, as the only basic amino acid. Statistical, chemical modification of ornithine sidechains to arginine promoted structure and function. The function concomitantly evolved – from phase separation with RNA (coacervates) to avid and specific dsDNA binding – thereby demonstrating a smooth, gradual peptide-to-protein transition with respect to sequence, structure, and function.

## Introduction

The very first, founding members of today’s protein families emerged *de novo*. Such *de novo* emergence is thought to begin with coincidental expression of polypeptides that have no prior physiological role^1^. If such polypeptides happen to provide some benefit, they may ultimately, through a series of duplications and fusions, and merging with other peptide fragments, yield a new folded and functional protein^2–5^. While also an ongoing process^6^, this mechanism of emergence foremost applies to the earliest proteins, regardless of whether the precursor polypeptides were formed by a primitive translation machinery, by other template-driven processes^7^, or by spontaneous assembly^8^. However, crossing the peptide-protein divide remains a poorly-understood evolutionary process. Specifically, an evolutionary trajectory leading from a primordial polypeptide to a modern protein regards three dimensions – sequence, structure, and function – that coevolve and need to be satisfactorily accounted for.

Changes in sequence regard an increase in length, but also the expansion of the amino acid alphabet. It is widely accepted that early proteins were predominantly comprised of “primordial amino acids” – *i.e.*, amino acids formed by spontaneous abiotic synthesis^9–12^. With time, this abiotic alphabet expanded to the twenty canonical amino acid set known today. It has also been postulated that the early proteins were statistical – that is, being the products of a primordial synthetic machinery, they comprised a mixture of related sequences^13,14^. Structure is also presumed to evolve from polypeptides with partial or complete disorder to ordered, tightly-packed globular domains ^4,15^. Finally, the refinement of sequence and structure allowed function to develop, for example, from ligand binding with low affinity and specificity to highly avid and selective binding.

Insights as to how a folded, biochemically-active protein can emerge from a simpler polypeptide are rare^16–18^, certainly when it comes to the natural protein world. Firstly, biochemical functions such as binding of small ligands demand a globular protein with a pre-organized active site. We thus lack knowledge as to what meaningful functions simple precursor polypeptides that lack a defined 3D structure could fulfill. Polypeptides, and even short peptides, with various biochemical functions have been described^16,19,20^, but the functions described so far have limited evolutionary relevance. Secondly, given a simple polypeptide with a rudimentary yet evolutionary-relevant function, how can this polypeptide gradually evolve to give a contemporary protein with a proficient and specific function?

Given the above questions, we attempted to construct a trajectory that starts with a simple polypeptide made of abiotic amino acids that would eventually lead to a contemporary folded domain. We aimed to demonstrate a smooth, gradual transition from a polypeptide to its modern protein descendant, with respect to sequence, structure, and function. Specifically, we explored the emergence of a nucleic acid-binding element. Early polypeptides, those emerging well before the last universal common ancestor (LUCA), likely interacted with polynucleic acids such as RNA^20–23^ and possibly also with DNA^24^. However, early emergence of nucleic acids binding presents a conundrum. Basic amino acids are critical for nucleic acid binding and must have been part of primordial proteins^20^. Yet none of the current proteogenic basic amino acids (Lys, Arg, His) are considered abiotic. Arginine is thought to be the earliest canonical basic amino acid^10,20^, and its precursor has been synthesized under abiotic conditions^25^. However, arginine is not found in simple abiotic syntheses nor in meteorites. Here, we propose a new hypothesis: ornithine, an abiotic amino acid that exists in today’s cells but not in today’s proteins, could have been the first basic amino acid. We further show that statistical chemical conversion of ornithine sidechains to arginine promotes both function and folding. Finally, we show that avid and selective dsDNA binding can emerge from a polypeptide that exerts a function that is presumed to be one of the earliest peptide-protein functions: formation of coacervates with RNA^26^.

## Results

### Our model case – the (HhH)_2_ fold

The helix-hairpin-helix (HhH) motif is a pre-LUCA nucleic acid-binding element found in at least eight protein superfamilies^27^, including ribosomal protein S13^28^, with broad distribution across the tree of life. Indeed, ribosomal proteins are likely the closest relics of the RNA-peptide world and of the earliest stages of evolution of folded proteins^21^. The HhH binding loop is simple and glycine-rich (consensus sequence: PGIGP) and can interact with both double- and single-stranded RNA and DNA^27^. HhH motifs can function as a single element within a larger domain or as a standalone 4-helix bundle, the (HhH)_2_ fold, in which two HhH motifs are symmetrically juxtaposed (**Figure 1a**). The pre-LUCA history, and the ability to form an independently-folded and functional protein, motivated us to select the (HhH)_2_ fold as the starting point for our study. We performed a systematic simplification of the (HhH)_2_ fold, akin to a retrosynthetic analysis in organic chemistry^29^, by generating experimental models of ancient evolutionary states.

**Figure 1.**
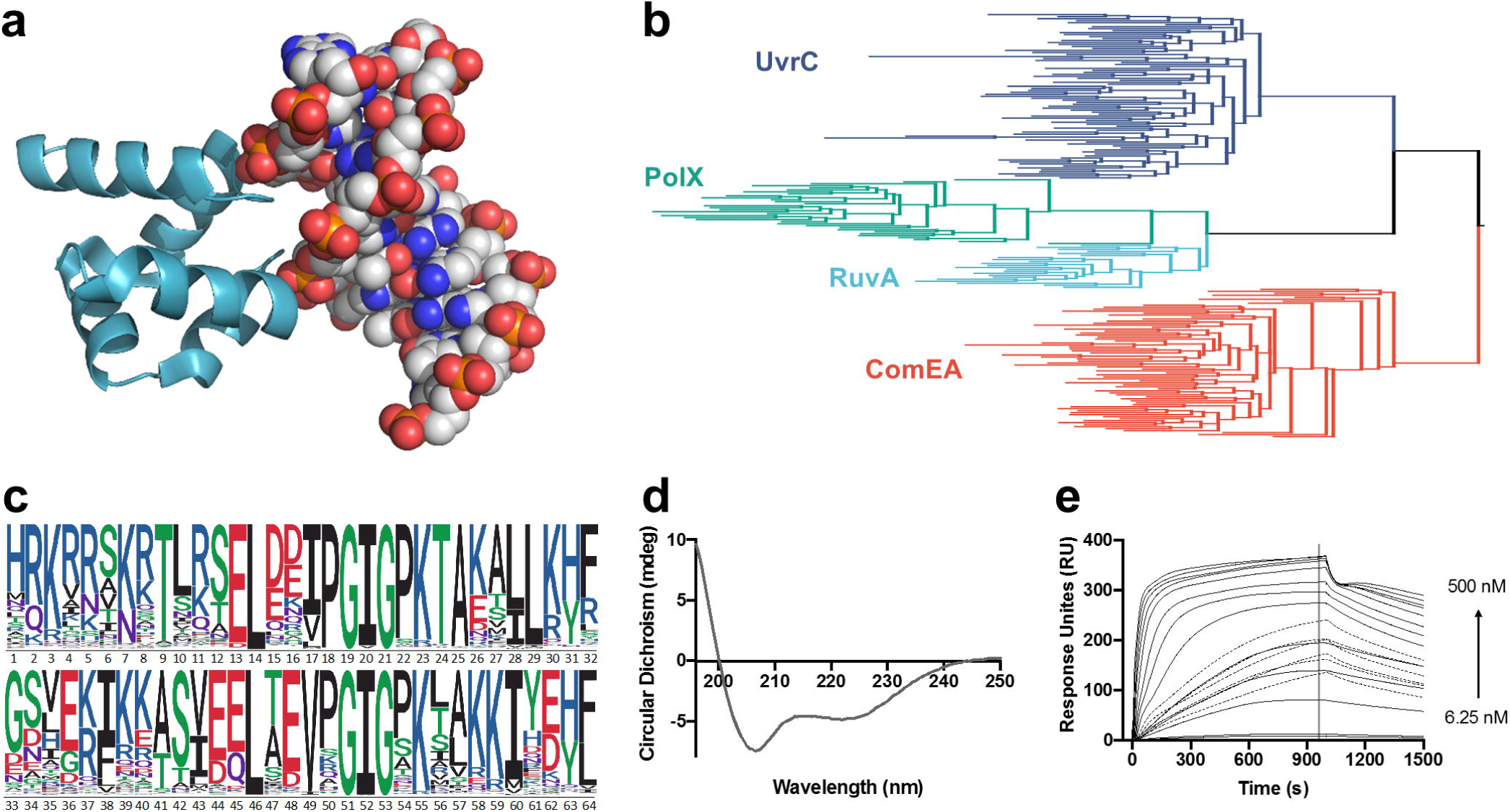
Reconstruction of the (HhH)_2_ ancestor. **a**. The characteristic (HhH)_2_ fold and its binding to the minor groove of dsDNA (shown is RuvA crystal structure, PDB code 1c7y). **b**. A midpoint-rooted tree of the four contemporary (HhH)_2_ protein families used for ancestor state inference. Crucially, sequences associated with the different (HhH)_2_ protein families segregated to monophyletic clades (blue, ComE; red, RuvA; green, PolX; orange, UvrC; sequences are listed in **Supplemental File 1**). **c**. A sequence logo representing the inferred ancestral amino acids for the common ancestor of the (HhH)_2_ lineage. Character height scales with posterior probability (the most probable sequence, dubbed Ancestor-(HhH)_2_, is listed in **Table 1**). The logo was modified to follow the topology of UvrC with respect to the linker connecting the two HhH motifs and other gaps. **d**. Circular dichroism (CD) spectrum of Ancestor-(HhH)_2_ showing the expected α-helical character. **e**. Binding of Ancestor-(HhH)_2_ to 101 base pair dsDNA fragment was measured by surface plasmon resonance (SPR). A synthesized version was tested on Biacore T200 (see Supplementary Methods). The association kinetics are biphasic, with fast initial binding followed by a second slower step (probably related to a structural rearrangement). At moderate concentrations there appears to be a change in the rate-limiting step (perhaps from initial binding being rate-limiting at low concentrations to structural isomerization becoming rate-limiting at the higher ones; see **Figure 3d** for plot of max RU). At these transition concentrations, even a contact time of 1000 s was insufficient to reach steady-state (dotted lines).

### Reconstruction of the last (HhH)_2_ common ancestor

Four contemporary (HhH)_2_ protein families are known – UvrC, PolX, RuvA, and ComEA – that comprise two fused HhH motifs and bind nucleic acids, usually dsDNA. A phylogenetic tree was constructed, indicating, as expected, four monophyletic clades that correspond to these four known families (**Figure 1b**). The last (HhH)_2_ common ancestor was then inferred using maximum likelihood methods (**Figure 1c**). For the core domain, the ancestor sequence was derived, as customary, from the most probable amino acid at each position^30^. However, the linker that connects the two HhH subdomains differs between (HhH)_2_ families in both sequence and length, and hence its ancestral state could not be reliably inferred. Although ComEA seems like the closest clade to the common (HhH)_2_ ancestor, its linker comprises an outlier with respect to the other three (HhH)_2_ families, and is also more complex, requiring additional N- and C-terminal segments to cap the hydrophobic core (**Figure S1a**). We thus opted for the UvrC linker, which is 8 residues long and also resembles the PolX and RuvA linkers. This linker was chosen due to its relative simplicity, and also because the last 4 residues of the linker (positions 37-40 in **Figure 1c**) happen to align well with the sequence of first α-helix (positions 5-8). This reduced the effective linker length to just 4 residues (positions 33-36) and further increased the symmetry of the ancestral sequence. The resulting protein, which represents the progenitor of the modern (HhH)_2_ families, is referred to as Ancestor-(HhH)_2_ (**Table 1**). Ancestor-(HhH)_2_, and other variants mentioned in the main text, were chemically synthesized to match their below-described ornithine version that could be made only by chemical synthesis (**Figure S1** panels **b**-**f**). This simple core domain was soluble, adopted the expected helical structure (**Figure 1d**), and avidly bound dsDNA (**Figure 1e**).

**Table 1.**
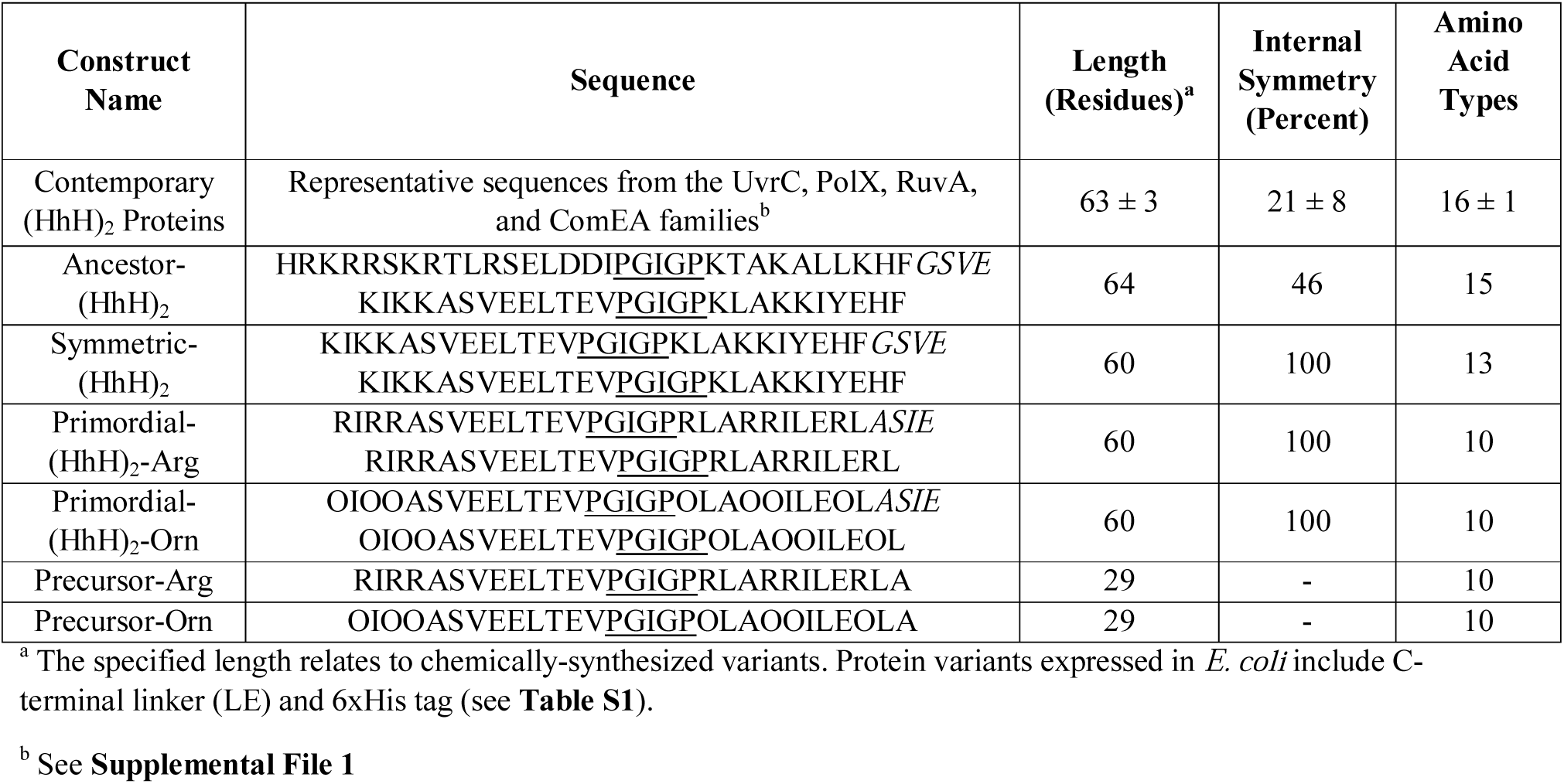
Sequences of HhH proteins and polypeptides.

Ancestor-(HhH)_2_ turned out to be significantly more symmetric than contemporary sequences, with 46% identity between the two HhH subdomains *versus* 21 ± 8% in extant (HhH)_2_ proteins (**Table 1**). Ancestral sequence reconstruction has no inherent bias for sequences with internal symmetry. Though the linker was also chosen to promote symmetry, this only increased the internal sequence identity by a single residue, or by 3.6%. The increased symmetry therefore supports the proposed scenario of emergence of the (HhH)_2_ domain *via* duplication and fusion of a polypeptide precursor^2^. To further support this mechanism of emergence, we attempted to construct an (HhH)_2_ intermediate with 100% sequence identity between the two HhH subdomains – *i.e.*, a functional protein where both subdomains have an identical sequence.

### A symmetrical (HhH)_2_ ancestor

Since traditional phylogenetic approaches do not enable reconstruction beyond the last common (HhH)_2_ ancestor, three alternative approaches were applied to achieve complete symmetrization (**Table S1**): First, we inferred the ancestral HhH motif from a tree comprising the first and second HhH subdomains of natural (HhH)_2_ proteins. The resulting sequence, however, when duplicated and joined by the UvrC linker, did not bind dsDNA.

Next, the first and second subdomains of Ancestor-(HhH)_2_ were duplicated while retaining the UvrC linker (and thereby symmetrizing positions 5-8 and 37-40; see above). Of these two variants, duplication of the second subdomain of Ancestor-(HhH)_2_ yielded a functional protein, dubbed Symmetric-(HhH)_2_ (**Table 1, Figure S2**). Finally, a third variant was constructed in which the most probable amino acid across the two symmetric halves was chosen. This third approach also yielded a functional protein (**Table S1**), indicating that multiple sequences can satisfy the symmetry constraint.

Symmetric-(HhH)_2_ has 100% sequence identity between subdomains, and is thus the conceptual product of duplication and fusion of a precursor polypeptide of 28 amino acids. It also utilizes only 13 amino acid types, compared to natural (HhH)_2_ proteins which use 16 ± 1 types (**Table 1**). Indeed, it is widely accepted that early proteins were predominantly comprised of “primordial amino acids” – *i.e.*, amino acids formed by spontaneous abiotic synthesis. Foremost, of the current 20 canonical amino acids, Gly, Ala, Ser, Thr, Asp, Glu, Pro, Val, Ile, and Leu, are considered abiotic^9–11^. Symmetric-(HhH)_2_, however, included several amino acids that likely emerged with enzyme-based biosynthesis, specifically His, Phe, Tyr, and Lys. Can a functional variant of the (HhH)_2_ fold be constructed that is both fully symmetric and comprised of an abiotic amino acid alphabet?

### A primordial (HhH)_2_ ancestor

Removal of His, Phe, and Tyr from Symmetric-(HhH)_2_ was guided by three considerations: (i) the second most probable amino acid from ancestor inference (**Figure 1c**), (ii) the consensus amino acid of the UvrC protein family, and (iii) abiotic amino acids with similar properties (*e.g.*, leucine or isoleucine instead of aromatic amino acids). In practice, these three approaches often yielded overlapping sequences (**Table S2**). Alternative core mutations based on the three considerations outlined above were also tested to ensure stable core packing. In all cases, 100% sequence identity between subdomains was preserved.

While phenylalanine and tyrosine were readily eliminated, the only viable substitution of histidine that we could identify was to lysine, which is not considered abiotic. Further, Symmetric-(HhH)_2_ already contained 12 lysine residues. We thus obtained a protein with 14 lysine residues that comprised >20% of its residues (14/60 residues), underscoring the crucial need for basic amino acids in nucleic acid binding^20^. Can these lysine residues be substituted to another, more ancient basic amino acid?

As arginine is thought to predate lysine^10,20^, we attempted to replace the lysine residues with arginine. We constructed a series of variants with increasing degrees of lysine to arginine exchange, ultimately replacing all 14 lysines to arginine. These exchanges were not only tolerated, but improved affinity for dsDNA (**Table S2**). The product of complete exchange, Primordial-(HhH)_2_-Arg (**Table 1**) adopted a helical structure and bound specifically to dsDNA (**Figure 2** and **Figure S3**). Structure and function were therefore achieved despite being constructed from only 10 different amino acid types – 9 primordial amino acid types plus arginine. NMR indicated that the Gly residues in the canonical PGIGP binding motif broaden upon addition of dsDNA, while residues near the Gly residues experienced progressive chemical shift changes as dsDNA was titrated in. This result suggests that the canonical PGIGP loops and neighboring residues mediate dsDNA binding (**Figure 2**), as further confirmed by deactivation mutations in the PGIGP loops of Symmetric-(HhH)_2_ (**Table S3**).

**Figure 2.**
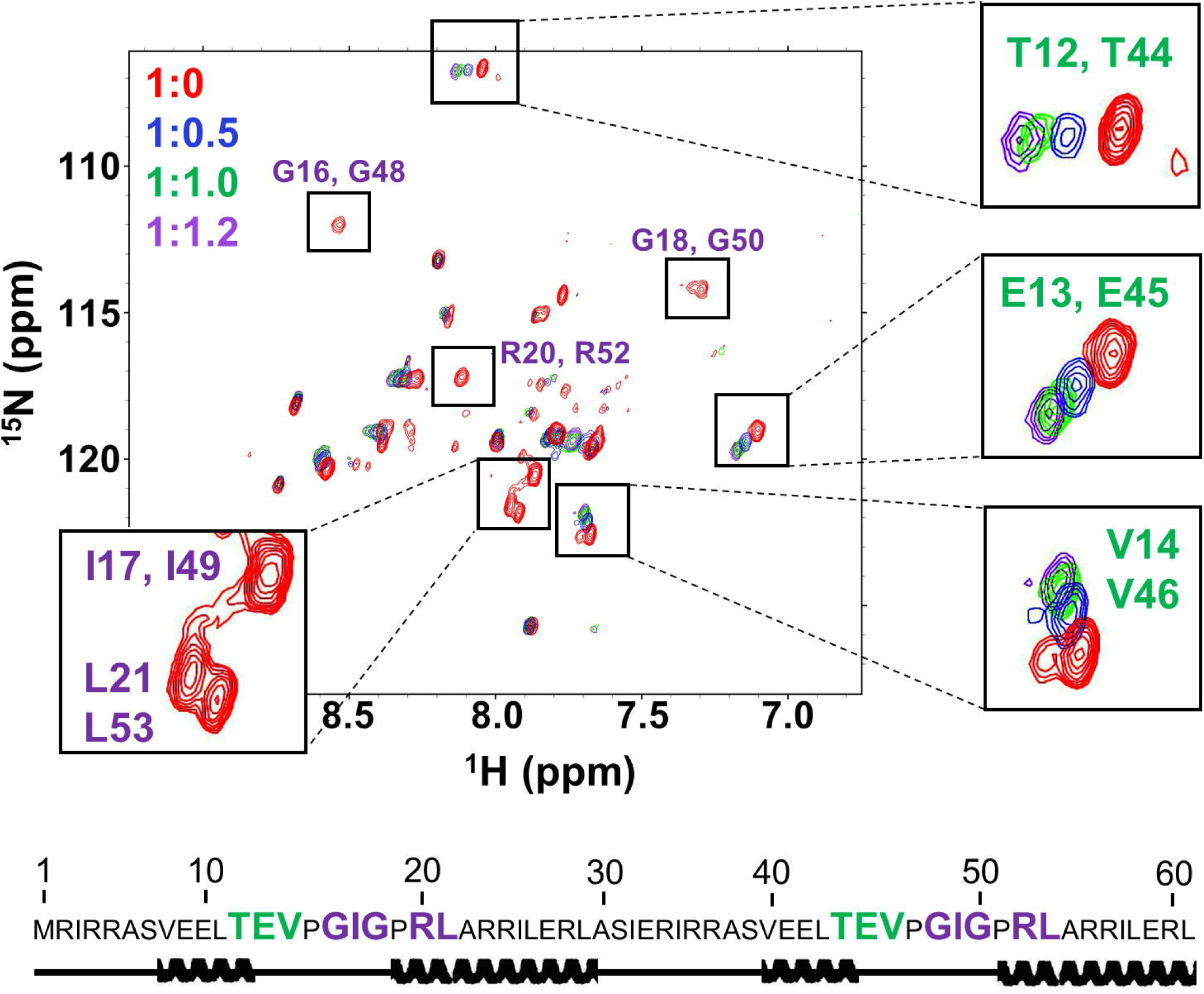
NMR characterization of Primordial-(HhH)_2_-Arg. ^1^H-^15^N HSQC titration of Primordial-(HhH)_2_-Arg with 12 base pair dsDNA. Protein:dsDNA ratios tested were 1:0 (red spectrum), 1:0.5 (blue spectrum), 1:1.0 (green spectrum), and 1:1.2 (magenta spectrum). Upon dsDNA binding, several peaks shifted due to fast exchange (green lettering) or broadened due to intermediate exchange regime (purple lettering). Notably, glycine peaks associated with the canonical binding motif PGIGP exhibited intermediate exchange binding behavior, suggesting the interaction with dsDNA is centered around the PGIGP motif. Secondary structure prediction was performed by TALOS+ using chemical shift alignments confirmed the helical structure of Primordial-(HhH)_2_-Arg (see also **Figure S3c**). For NMR studies, *E. coli* expressed tag-free Primordial-(HhH)_2_-Arg was used. Numbering is based on the *E. coli* expressed protein, in which residue 1 is methionine. A 12 base pair dsDNA oligonucleotide was used for NMR experiments to minimize avidity effects in binding and retain fast molecular tumbling during dsDNA titration. The DNA sequences for all experiments can be found in **Table S5**.

Overall, Primordial-(HhH)_2_-Arg differs from the last common (HhH)_2_ ancestor at 32 out of 64 positions, including 12 lysine and 2 histidine residues replaced by arginine. When compared to modern (HhH)_2_, and considering only positions that align to the last common (HhH)_2_ ancestor, Primordial-(HhH)_2_-Arg differs at 45 ± 4 out of 64 positions. Nonetheless, its folding and binding mode are consistent with those of contemporary (HhH)_2_ domains.

### A functional ornithine-based (HhH)_2_ ancestor

Arginine likely emerged before lysine and histidine, yet it is still not considered among the earliest, abiotic amino acids. This led us to examine whether the (HhH)_2_ fold could possibly function with another basic amino acid that could have predated arginine. Ornithine is the only basic amino acid so far identified in a volcanic spark discharge experiment^11^ and in meteorites^31^ and is a metabolic precursor of arginine^32^. Could the first basic amino acid be ornithine?

Since ornithine cannot be incorporated by standard protein expression approaches, we resorted to chemical protein synthesis. Two half-peptides were synthesized and then joined using a native chemical ligation (NCL) and a deselenization approach^33,34^. To permit deselenization chemical approaches, the glycine in the linker connecting the two HhH motifs of Primordial-(HhH)_2_ was changed to alanine (with no effect on folding or binding, as tested on the arginine version; **Table S2**). For each construction, the N-terminal half-peptide was synthesized and its C-terminal thioester was reacted with the C-terminal peptide that had an N-terminal selenocysteine. Following ligation of the two peptides, the selenocysteine residue was deselenized to yield alanine (see **Methods**).

Remarkably, Primordial-(HhH)_2_-Orn (**Table 1, Figure S4**), in which all 14 arginine residues have been replaced by ornithine, weakly bound dsDNA (**Figure S4f** and **Figure 3c**), indicating that, in principle, functional dsDNA binding proteins can be produced in the absence of either arginine or lysine. Circular dichroism spectroscopy indicated that Primordial-(HhH)_2_-Orn is largely unfolded, though addition of dsDNA seems to induce some helical structure (**Figure S4g**). Likewise, exchanging lysine for ornithine in the context of Symmetric-(HhH)_2_ also yielded a functional protein (**Figure S5**).

**Figure 3.**
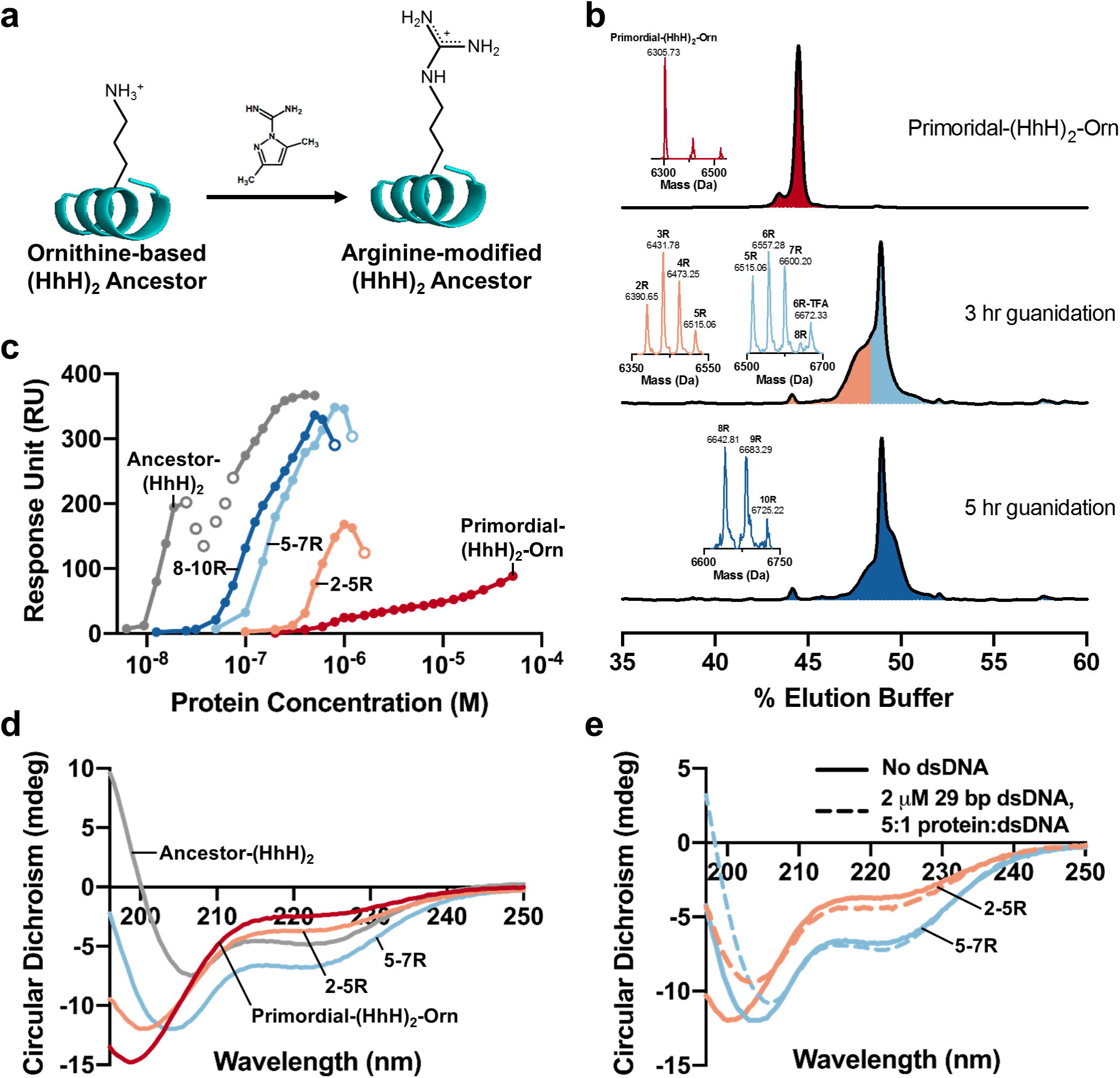
Statistical conversion of ornithine sidechains to arginine promotes structure and dsDNA-binding. **a**. A simple chemical reaction, guanidination, performed in water can convert the ornithine sidechains of a chemically synthesized protein into arginine sidechains (see also **Figure S6** and **Figure S7**). **b**. The synthesized Primordial-(HhH)_2_-Orn analyzed by reverse-phase chromatography and mass spectrometry (expected and measured masses are listed below; top chromatogram). Three mixtures of Primordial-(HhH)_2_-Orn with different degrees of guanidination of its 14 ornithine residues were prepared by varying the reaction time and fractionation by reverse-phase chromatography (middle and bottom chromatograms). These mixtures contained 2-5 guanidinations per polypeptide (orange), 5-7 guanidinations per polypeptide (light blue), and 7-10 guanidinations per polypeptide (dark blue). The composition of these mixture was determined by mass spectrometry analysis. Note that a single mass represents a mixture of proteins guanidinylated to the same degree yet at different positions. **c**. The binding affinity for dsDNA increases monotonically with the extent of guanidination, as indicated by surface plasmon resonance binding experiments with an immobilized 101 base pair dsDNA fragment (see **Figure S8d** for an equivalent plot of binding to 29 base pair dsDNA). Open circles indicate concentrations where a change in rate-limiting step occurred (see **Figure 1e** for additional details). **d**. The α-helical character of the protein increases with increasing guanidination, as demonstrated by circular dichroism spectroscopy (195 nm signal becomes increasingly positive, the signal at 222 nm increasingly negative, and the global minimum shifts towards 208 nm). **e**. An increase in α-helicity is observed upon mixing protein with 29 base pair dsDNA, suggesting folding upon binding. Plotted spectra are background-subtracted to remove the signal associated with dsDNA. Mass spectrometry confirmation: Primordial-(HhH)_2_-Orn (m_calc_ = 6307.57 Da, m_obs_ = 6305.73 ± 0.56 Da); Primordial-(HhH)_2_-2R (m_calc_ = 6391.65 Da, m_obs_ = 6390.65 ± 2.02 Da); Primordial-(HhH)_2_-3R (m_calc_ = 6433.69 Da, m_obs_ = 6431.78 ± 0.30 Da); Primordial-(HhH)_2_-4R (m_calc_ = 6475.73 Da, m_obs_ = 6473.25 ± 0.99 Da); Primordial-(HhH)_2_-5R (m_calc_ = 6517.77 Da, m_obs_ = 6515.06 ± 0.81 Da); Primordial-(HhH)_2_-6R (m_calc_ = 6559.81 Da, m_obs_ = 6557.28 ± 1.80 Da); Primordial-(HhH)_2_-6R-TFA (m_calc_ = 6673.83 Da, m_obs_ = 6672.33 ± 3.66 Da); Primordial-(HhH)_2_-7R (m_calc_ = 6601.85 Da, m_obs_ = 6600.20 ± 1.43 Da); Primordial-(HhH)_2_-8R (m_calc_ = 6643.89 Da, m_obs_ = 6642.81 ± 1.00 Da); Primordial-(HhH)_2_-9R (m_calc_ = 6685.93 Da, m_obs_ = 6683.29 ± 1.32 Da); Primordial-(HhH)_2_-10R (m_calc_ = 6727.97 Da, m_obs_ = 6725.22 ± 2.12 Da).

### Statistical conversion to arginine promotes folding and dsDNA binding

Ornithine has another distinctly attractive feature. In contemporary metabolism, ornithine, *i.e.*, the free amino acid, is guanidinated by a series of enzymatic reactions to give arginine that is, in turn, incorporated into proteins^32^. However, guanidination of ornithine to give arginine can also be performed chemically, and even by simple reagents such as cyanamide (H_2_N-CN)^35^, which are likely to have been present in primordial environments. Foremost, chemical guanidination can be readily performed on polypeptides that contain ornithine, as a post-synthesis modification that leads to arginine (**Figure 3a**). This reaction, which operates on already-synthesized peptides, was reported^35^, but its products were not fully characterized. We tested this protocol on model polypeptides and observed complete conversion of ornithine into arginine in aqueous solution. Conversion was also selective, as no modification of the peptide’s N-terminal amine group, or a histidine side chain, were observed (**Figure S6** and **Figure S7**).

The reaction was then applied to Primordial-(HhH)_2_-Orn to achieve partial conversion (**Figure 3b**). Guanidinated mixtures of Primordial-(HhH)_2_-Orn were then isolated. Although these mixtures were random with respect to which ornithine positions were converted to arginine, they had discrete compositions with respect to the total number of modified ornithine residues (**Figure 3b**). We found that greater conversion of ornithine to arginine resulted in progressive improvement in both the affinity for dsDNA and folding towards the expected (HhH)_2_ helical structure, as judged by SPR (**Figure 3c** and **Figure S8)** and circular dichroism (**Figure 3d** and **3e**), respectively.

### A single, primordial HhH motif forms coacervates with RNA

The results described above show that the (HhH)_2_ fold had likely emerged from duplication of a short, simple polypeptide comprised of just one HhH motif. In principle, the precursor polypeptide could emerge by chance and, once duplicated, become evolutionarily advantageous. However, emergence from an already functional single HhH motif polypeptide is far more likely, as nascent activity is a viable starting point for evolutionary optimization^3,18^. We thus tested pre-duplicated 29-residue polypeptides derived from Primordial-(HhH)_2_, containing either ornithine (“Precursor-Orn”) or arginine (“Precursor-Arg”) (**Table 1**). Albeit, none of these polypeptides showed reproducible dsDNA binding.

The functional and structural properties of ancient polypeptides have been a matter of intense interest. In principle, binding small molecule ligands, not to mention catalysis, demands structural volume and complexity, and a degree of pre-organized structure, that polypeptides rarely provide. It has thus been postulated, however, that self-assembly could endow primordial polypeptides such properties. Specifically, both amyloid formation^36^ and peptide-RNA condensates^26^ have been postulated as ancient forms of self-assembly. Could formation of liquid condensates with RNA be an early function that pre-dated nucleic acid binding in modern proteins?

Upon mixing with polyuridylic acid (PolyU), both Precursor-Orn and -Arg formed liquid coacervates (**Figure 4a** and **Figure S9**), as did Ancestor-(HhH)_2_ (**Figure S10**). Neither polypeptide formed droplets in the absence of PolyU. Further, precursor-Arg consistently made larger droplets than Precursor-Orn, and at lower concentrations (**Figure S11**). Thus, by increasing the coacervate-forming potential, statistical modification of ornithine to side chains to arginine could also provide an advantage at this early stage.

**Figure 4.**
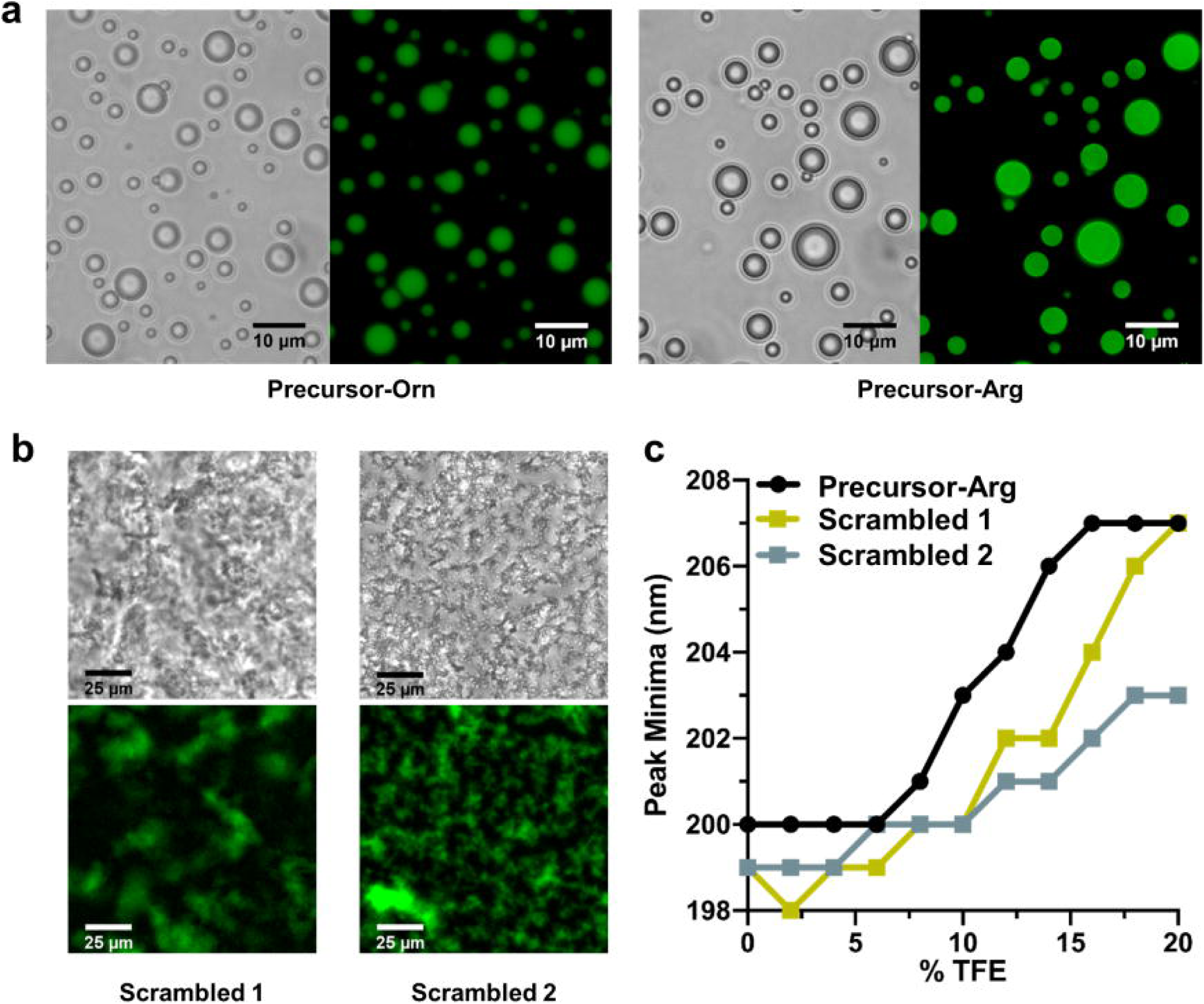
Coacervates formation by Precursor polypeptides and RNA. **a**. Coacervates form upon addition of poly-uridylic acid (polyU) to both Precursor-Orn (240 μM peptide, 1.2 μM fluorescein-labeled peptide, 1.2 mg/ml polyU) and Precursor-Arg (190 μM peptide, 20 μM fluorescein-labeled peptide, 1.4 mg/ml polyU). **b**. Complete scrambling of the sequence of Precursor-Arg (“Scrambled 1”), as well as exclusion of the arginine residues (“Scrambled 2”), abolishes droplet formation (200 μM peptide, 8 μM fluorescein-labeled peptide, 1 mg/ml polyU; see **Table S4** for polypeptide sequences). **c**. Triflouroethanol (TFE) titrations of the polypeptides in buffer reveal that Precursor-Arg has a higher intrinsic α-helical propensity than the scrambled versions. This is demonstrated by a shift in the circular dichroism peak minimum from ∼199 nm (random coil-like) to 207 (α-helix-like) at lower concentrations of TFE (10 μM polypeptide; see for **Figure S12** for raw CD spectra).

Phase separation of these polypeptides is, perhaps, unsurprising given that arginine-rich peptides have been shown to phase separate with addition of RNA or even crowding reagents^37^. However, scrambling the sequence of the Precursor-Arg – either completely, or while preserving only the positions of the arginine residues (scrambled 1 and 2, respectively) – resulted in amorphous aggregates upon PolyU addition (**Figure 4b**). Thus, basic amino acids are likely necessary, but not sufficient, to encode phase separation.

Finally, triflouroethanol titration experiments with the polypeptides in solution demonstrated that the Precursor-Arg that corresponds to a single HhH element has, as expected, a higher propensity to form α-helices than the scrambled variants (**Figure 4c** and **Figure S12**). This suggests that sequences that evolved to phase separate may have a higher propensity to fold into a helical structure upon duplication.

## Discussion

Proteins with defined biochemical function are thought to be exceedingly rare in sequence space, with some probability estimates as low as 10^−11^ for nucleotide binding^38^. However, the trajectory described here suggests a possible solution to this evolutionary conundrum, whereby evolution begins with a rudimentary function, such as phase separation, that can be mediated by polypeptides with minimal sequence constrains. Duplication events, and expansion of the amino acid alphabet, would then enable a smooth transition to more complex protein forms, with both structure and function becoming increasingly defined over time. During simplification, ∼70% of all positions in Ancestor-(HhH)_2_ were changed to generate Primordial-(HhH)_2_-Arg. These changes included 14 simultaneous exchanges of the basic amino acids that were tolerated, also to the currently non-proteogenic ornithine, while maintaining dsDNA binding. There is therefore a large cloud of traversable sequences from which the actual, historical ancestor of the HhH element could have emerged. This unusual tolerance to sequence variation, and emergence by duplication, indicate a probability of emergence much higher than 10^−11^.

Our results also show that polypeptides and proteins can bind polynucleic acids with ornithine as the sole basic amino acid. Nucleic acid binding is a primordial function that likely emerged in a peptide-RNA world^8,20,21^. We thus offer a new hypothesis by which ornithine was the main basic amino acid in the earlies stages of protein evolution. The evolutionary change in amino acid composition and the genetic code is often viewed as going from a subset of the current twenty proteogenic amino acids to the complete set. However, many other amino acids are seen in abiotic synthesis experiments and/or meteorites, including amino acids that are ubiquitous in modern biology yet are not incorporated into proteins (nonproteogenic). Some of these amino acids are thought to have been included in early proteins, such as norvaline and α-aminobutyric acid^39^. Indeed, the earliest hypotheses regarding the emergence of translation and the genetic code assumes a large primordial set of amino acids from which the current set of 20 canonical amino acids was selected^13,14^. Ornithine is present in living cells but not encoded by ribosomal protein synthesis, the primary explanation being the presumed instability of its tRNA ester (due to intramolecular lactam formation^40^). However, that ornithine is present in nonribosomal peptides^41^ indicates that its esters (and the even more reactive thioesters) are sufficiently long-lived to allow its condensation into polypeptides. Foremost, ornithine is the biosynthetic precursor of arginine. There are multiple indications that intermediates of contemporary metabolic pathways served as end-point metabolites at earlier evolutionary stages^32^. Thus, arginine may have first emerged as a spontaneous, chemical modification of ornithine sidechains in already synthesized polypeptides. The superior properties of arginine, in promoting coacervate formation, as well as dsDNA binding and folding, and the instability of ornithine esters, led to the takeover of arginine as a protein building block ^23^.

Finally, this experimentally-validated trajectory, which starts from a polypeptide that forms coacervates with RNA and leads to a contemporary RNA/DNA binding protein, lends support to a hypothesis that stems from Oparin’s protocells hypothesis – namely, that peptide-RNA condensates have had a role in the earliest life forms^26^.

## Acknowledgements

We thank Brian Ross and Vladimir Kiss for assistance with fluorescence microscopy, Dora Tang for assistance with droplet imaging, Einav Aharon for assistance with molecular cloning, Jelena Cveticanin for assistance with mass spectrometry, Reem Mousa for assistance with polypeptide guanidination, and Ita Gruić-Sovulj for suggesting the use of ornithine as an alternative basic amino acid. We acknowledge insightful comments on this manuscript from Andrei Lupas, Donald Hilvert, and James Bardwell. This work was funded by the Israel Science Foundation (grant 980/14 awarded to D.S.T. and grant 783/18 awarded to N.M.), an ICRF Acceleration Grant (awarded to N.M.), and the National Institute of Health (grant R35 GM126942 awarded to G.V.). D.S.T. is the Nella and Leon Benoziyo Professor of Biochemistry. O.W.K. acknowledges the Kaete Klausner Fellowship for financial support.

