## Supplementary Material for "Primordial emergence of a nucleic acid binding protein via phase separation and statistical ornithine to arginine conversion"

### Supplementary Tables

| Construct Name | Sequence | ds/ssDNA Binding | Design Notes |
|---|---|---|---|
| HhH-1 Duplicated | M**RSKRTLRSELDDIPGIGPKTAKAL LKHF**GSVE**RSKRTLRSELDDIPGIGP KTAKALLKHF**LEHHHHHH | -/nd | Duplication of the first HhH subdomain of Ancestor-(HhH)$_2$. |
| HhH-2 Duplicated<br>**Symmetric-(HhH)$_2$** | M**KIKKASVEELTEVPGIGPKLAKKI YEHF**GSVE**KIKKASVEELTEVPGIGP KLAKKIYEHF**LEHHHHHH | ++/- | Duplication of the second HhH subdomain of Ancestor-(HhH)$_2$. |
| Sym100_short | M**TSVEELTEVPGIGPKTAKKILKHF** GSVEKIKK**TSVEELTEVPGIGPKTAK KILKHF**LEHHHHHH | ++/- | Most probable amino acid at each position after summing the posterior probabilities of the aligned sites from the ancestral sequence reconstruction. The 4-residue N-terminal poly-basic tail was also truncated. |
| Ancestor of (HhH)$_2$ Halves | M**NIKKASLEELAKVPGIGPKTAKKI YDYL**GSVE**NIKKASLEELAKVPGIG PKTAKKIYDYL**LEHHHHHH | -/nd | Ancestor of the HhH subdomains derived from an alignment of the HhH subdomain sequences. |

**Table S1. Symmetrization of Ancestor-(HhH)$_2$.** In all constructs summarized in the table, the UvrC linker (underlined) was used to connect two identical HhH subdomains (bold). Note that for chemically-synthesized proteins, ASVE was used as the linker sequence instead. Binding to DNA was assessed by ELISA (see **Materials and Methods**) and is classified as strong (+++), moderate (++), weak (+), undetectable (-), or not determined (nd).

| Construct Name | Soluble Expression Relative to Symmetric-(HhH)$_2$ | ds/ssDNA Binding | Design Notes |
|---|---|---|---|
| Symmetric-(HhH)$_2$ G29A | Same | ++/nd | Linker mutation to simplify total protein synthesis. |
| Symmetric-(HhH)$_2$ F28L/F60L | Increased | +/nd | Phe replaced to Leu which is the second most probable amino acid at position 60 in the inferred ancestor. |
| Symmetric-(HhH)$_2$ H27A/H59A | Decreased | -/nd | Substitution of His to the simplest residue with high helical propensity. |
| Symmetric-(HhH)$_2$ H27S/H59S | Decreased | -/nd | Substitution of His to the simplest polar residue. |
| Symmetric-(HhH)$_2$ H27E/F28L/H59E/F60L | Same | -/nd | Substitution of His to Glu, the second most common residue at position 59 in the consensus sequence. |
| Symmetric-(HhH)$_2$ H27K/H59K | Same | ++/- | Substitution of His with another basic amino acid. |
| Symmetric-(HhH)$_2$ H27R/H59R | Same | ++/nd | Substitution of His with another basic amino acid. |
| Symmetric-(HhH)$_2$ K1R/K3R/K4R/K33R/K35R/K36R | Decreased | ++/nd | Six of the 12 Lys residues converted to Arg. |
| Symmetric-(HhH)$_2$ K1R/K3R/K4R/K22R/K23R/K33R/K35R/K36R/K54R/K55R | Decreased | +++/nd | Ten of the 12 Lys residues converted to Arg. |
| Symmetric-(HhH)$_2$ K1R/K3R/K4R/K19R/K22R/K23R/K33R/K35R/K36R/K51R/K54R/K55R [Symmetric-(HhH)$_2$-Arg] | Decreased | +++/- | Complete exchange of Lys to Arg. |
| Symmetric-(HhH)$_2$-Arg V13I/Y25L/H27R/F28L/G29A/V45I/Y57L/H59R/F60L [**Primordial-(HhH)$_2$-Arg Version 1**] | Decreased | +/- | Primordial variant with alternative core packing (V13I and V45I). Combines the above simplification mutations, as well as exchange of Tyr to Leu which is the most probable amino acid at position 25 in the inferred ancestor. |
| Primordial-(HhH)$_2$-Arg Version 1 I13V/I45V | Decreased | +/- | Reversion of V13I and V45I mutations |
| Symmetric-(HhH)$_2$-Arg Version 1 I13V/V31I/I45V [**Primordial-(HhH)$_2$-Arg**] | Decreased | +/- | Primordial-(HhH)$_2$-Arg Version 1 plus linker optimization, to give the final primordial design (sequence listed in **Figure 1**). |

**Table S2: Alphabet simplification of Symmetric-(HhH)$_2$.** Starting with Symmetric-(HhH)$_2$ (**Table S1**), the non-prebiotic amino acids (Phe, His, Tyr, and Lys) were systematically replaced with prebiotic amino acids. Mutations were made simultaneously at both symmetry-related sites. Numbering is based on the synthetic constructs (*i.e.*, without methionine as the first residue). Binding to DNA was assessed by ELISA (see **Materials and Methods**) and is classified as strong (+++), moderate (++), weak (+), undetectable (-), or not determined (nd).

| Construct Name | Soluble Expression Relative to Parent | dsDNA Binding | Design Notes |
|---|---|---|---|
| Symmetric-$(HhH)_2$ I16K/I48K | Increased | - | Aliphatic to charged mutation to deform the binding loop. |
| Symmetric-$(HhH)_2$ G17E/G49E | Increased | - | Mutation of conserved Gly to a larger acidic amino acid to block the binding surface. |
| Symmetric-$(HhH)_2$ G17Q/G49Q | Increased | - | Mutation of conserved Gly to a larger polar amino acid to block the binding surface. |
| Primordial-$(HhH)_2$-Arg I16K/I48K | No expression | nd | Aliphatic to charged mutation to deform the binding loop. |
| Primordial-$(HhH)_2$-Arg G17E/G49E | No expression | nd | Mutation of conserved Gly to a larger charged amino acid to block the binding surface. |
| Primordial-$(HhH)_2$-Arg G17Q/G49Q | No expression | nd | Mutation of conserved Gly to a larger polar amino acid to block the binding surface. |
| Primordial-$(HhH)_2$-Arg Version B G17E/G49E | Increased | - | Mutation of conserved Gly to a larger charged amino acid to block the binding surface. |

**Table S3: HhH motif inactivation mutants.** To confirm that dsDNA binding is mediated by the PGIGP binding loop characteristic of the HhH motif, several loop mutations were tested. Numbering is based on the synthetic constructs (*i.e.*, without methionine as the first residue). Binding to dsDNA was assessed by ELISA (see **Materials and Methods**) and is classified as strong (+++), moderate (++), weak (+), undetectable (-), or not determined (nd). The I-to-K mutations were introduced to exclude the possibility of binding by non-specific electrostatic interactions. Due to the tightly folded nature of the hairpin motif, half of the mutations destabilized the protein resulting in loss of expression.

| Construct Name | Sequence | Design Notes |
|---|---|---|
| Precursor-Arg | RIRRASVEELTEVPGIGPRLARRILERLA | All basic residues are Arg (29 residues total length) |
| Precursor-Orn | OIOOASVEELTEVPGIGPOLAOOILEOLA | All basic residues are Orn (29 residues total length) |
| Precursor-Arg, All Scrambled | GEPAIRSEPIRLRLEGVRALRTEVGLRIRA | All positions of Primordial-HhH Arg scrambled (30 residues). |
| Precursor-Arg, Scrambled Except R | GRVRRSEIPLGIELEAGTLRLARRPEIRVA | All positions except Arg in Primordial-HhH-Arg scrambled (30 residues). |

**Table S4: Synthetic polypeptides.** In each case, a C-terminal Ala was included to enable ligation to generate the $(HhH)_2$ variants. For the scrambled variants, an N-terminal Gly was added to enable labeling with a fluorescent dye. The amino acid ornithine is denoted by the single letter code O. All peptides except Precursor-Orn were custom synthesized by Synpeptide Co. Ltd (see **Materials and Methods**).

| Type | Experiment(s) | Length | Forward Sequence |
|---|---|---|---|
| dsDNA | SPR, ELISA | 101 bp | 5'biotin-CCGTCCGTAATCATGGTCATAGCTGTTTCGT TTAAAATGAAGATACGGCGCGATGATACGCGTCGG GTTGTCTCTCTGTTGATACAGAGATACTAGATGTA-3' |
| dsDNA | SPR, ELISA | 29 bp | 5'biotin-CCGTCCGTAATCATGGTCATAGCTGTTTC-3' |
| dsDNA | CD titration | 29 bp | 5'-CCGTCCGTAATCATGGTCATAGCTGTTTC-3' |
| dsDNA | NMR titration | 12 bp | 5'-TAGATCGATCGC-3' |
| ssDNA | ELISA | 29 bases | 5'biotin-CCGTCCGTAATCATGGTCATAGCTGTTTC-3' |

**Table S5: DNA sequences used for the binding assays**.

#### Supplementary Figures

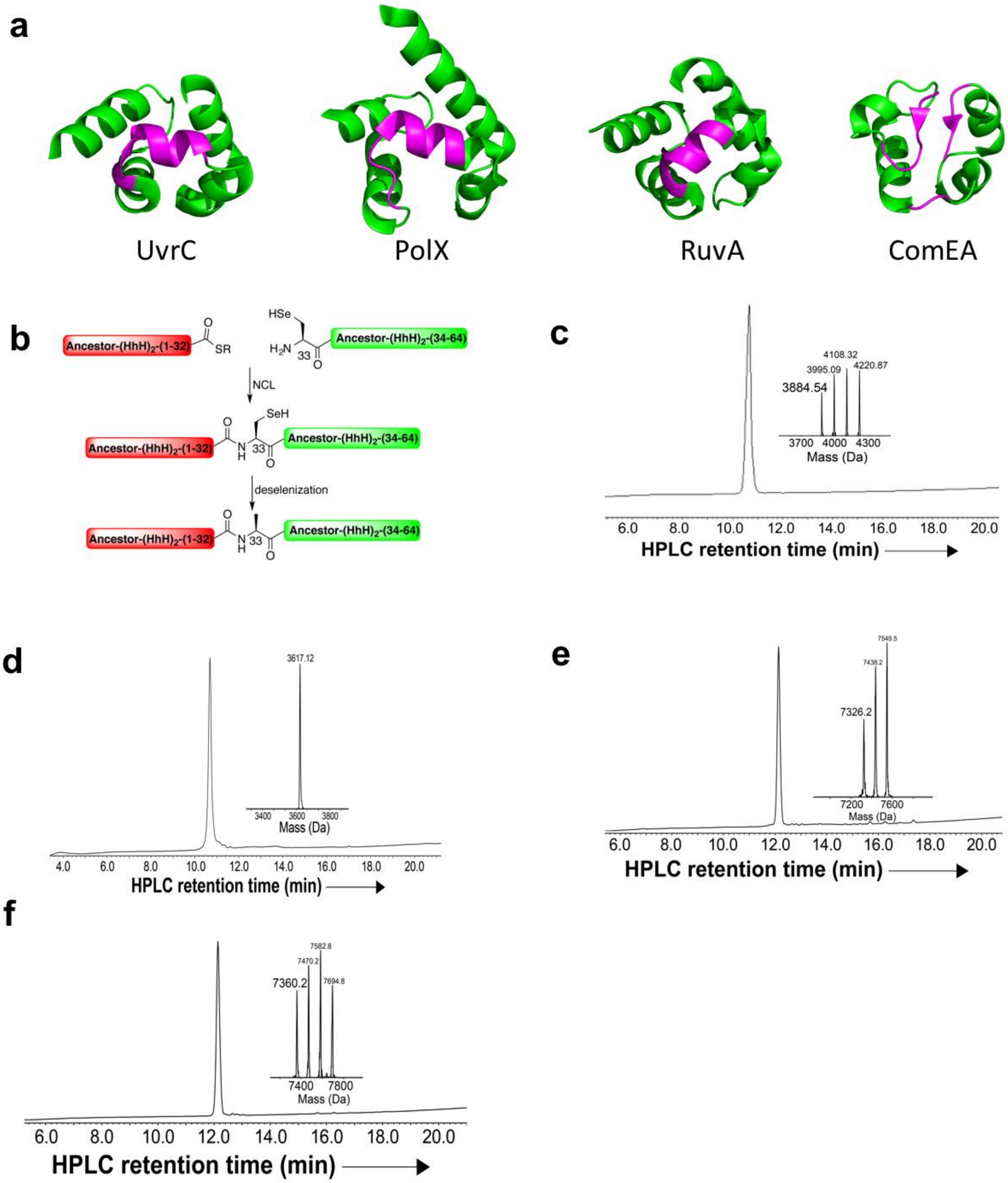


**Figure S1. Reconstruction and synthesis of Ancestor-(HhH)$_2$. a**. The different linker regions of the contemporary (HhH)$_2$ families. Three out of these four families, UvrC, PolX, and RuvA, use a short α-helix to link the two HhH subdomains and also cap the hydrophobic core (PDB accession codes 2nrt, 3b0x, and 2ztc, respectively). ComEA (PDB accession code 2duy), however, caps the hydrophobic core with a short intervening loop that connects the HhH subdomains and N- and C-terminal extensions that adopt a short β-sheet-like conformation. The linker for Ancestor-$(HhH)_2$ was chosen to be UvrC-like because of the latter's dominance among the contemporary families and its simplicity. Additionally, assigning the UvrC linker increased the internal sequence similarity of the inferred ancestor, thereby promoting further simplification (see main text for more details). **b**. Chemical protein synthesis scheme for Ancestor-$(HhH)_2$. Because the protein was too long for a single solid phase synthesis reaction, two half-peptides were synthesized and then joined using a native chemical ligation (NCL) and a deselenization approach. The N-terminal half-peptide bears the C-terminal thioester surrogate *N*-acyl urea, Nbz[1] moiety (NHalf-COSR) and the C-terminal peptide bears an N-terminal selenocysteine (Sec, U) residue (Sec-CHalf). After peptide ligation was complete, the selenocysteine residue was deselenized to yield alanine. **c**. HPLC chromatogram and mass spectrum demonstrating successful synthesis of Ancestor-$(HhH)_2$-NHalf-COSR ($m_{calc}$ = 3885.56 Da, $m_{obs}$ = 3884.54 Da, 3995.09 Da [M+TFA], 4108.32 Da [M+2TFA], and 4220.87 Da [M+3TFA]). **d**. HPLC chromatogram and mass spectrum demonstrating successful synthesis of fragment Ancestor-$(HhH)_2$-Sec-CHalf ($m_{calc}$ = 3619.13 Da, $m_{obs}$ = 3617.12 Da). **e**. HPLC chromatogram and mass spectrum demonstrating successful NCL of the half peptides to yield Ancestor-$(HhH)_2$-(A33U), in which residue 33 is selenocysteine ($m_{calc}$ = 7327.52 Da, $m_{obs}$ = 7326.2 Da, 7438.2 Da [M+TFA], 7549.5 Da [M+2TFA]). **f**. HPLC chromatogram and mass spectrum demonstrating successful deselenization of Ancestor-$(HhH)_2$-(A33U) to yield Ancestor-$(HhH)_2$ ($m_{calc}$ = 7361.6 Da, $m_{obs}$ = 7360.2 Da, 7470.2 Da [M+2TFA], 7582.8 Da [M+3TFA], 7694.8 Da [M+4TFA]).

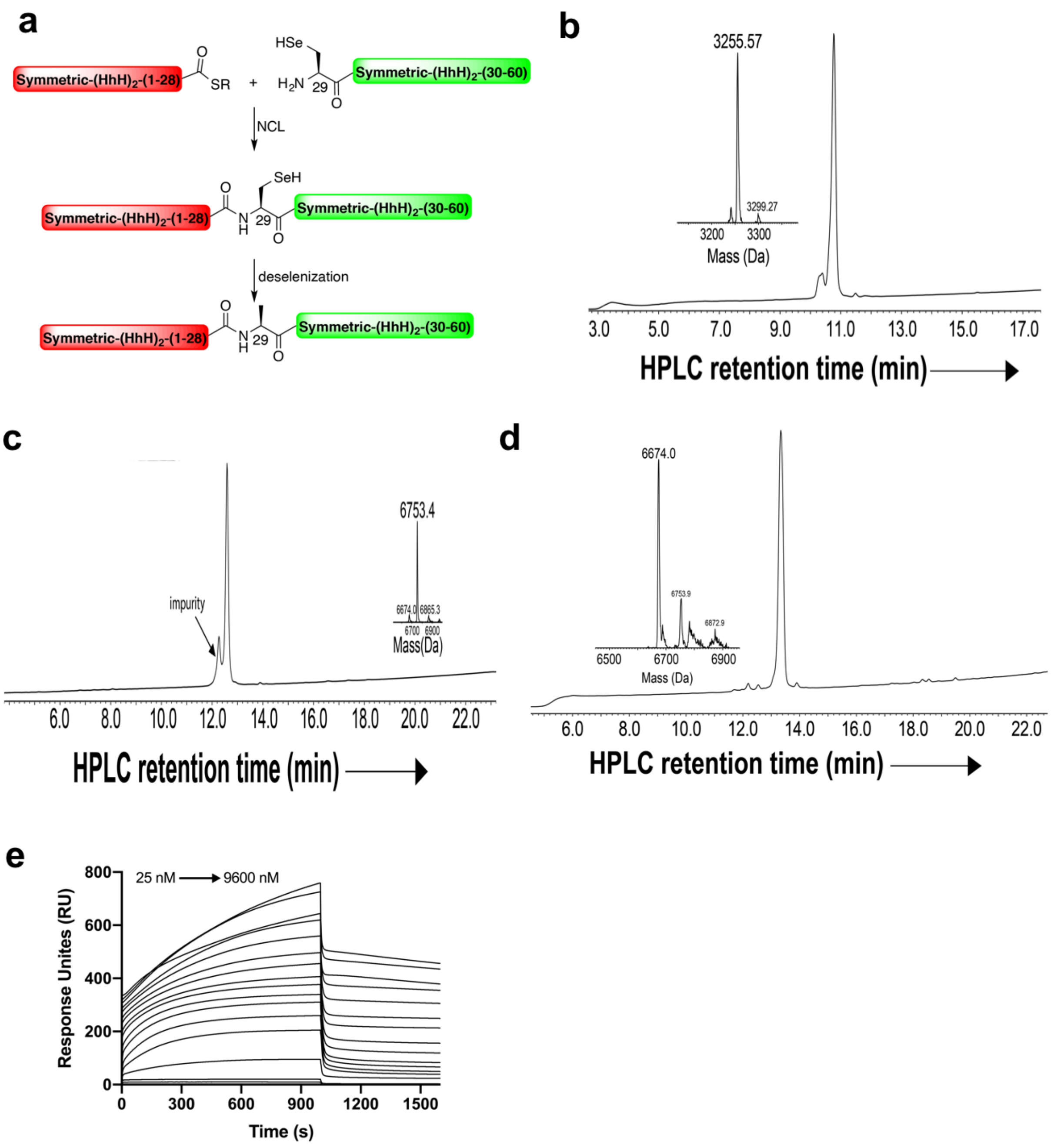


**Figure S2. Synthesis and characterization of Symmetric-(HhH)$_2$. a**. Chemical protein synthesis scheme for Symmetric-(HhH)$_2$. Because the protein was too long for a single solid phase synthesis reaction, two half-peptides were synthesized and then joined using an NCL and deselenization approach. The N-terminal half-peptide bears the C-terminal thioester surrogate *N*-acyl urea, Nbz moiety (NHalf-COSR), and the C-terminal peptide bears an N-terminal selenocysteine (Sec, U) residue (Sec-CHalf). After peptide ligation, the selenocysteine residue is deselenized to yield alanine. **b**. HPLC chromatogram and mass spectrum demonstrating successful synthesis of Symmetric-(HhH)$_2$-NHalf-COSR ($m_{calc}$ = 3255.92 Da, $m_{obs}$ = 3255.57 Da). **c**. Note that Symmetric-(HhH)$_2$-Sec-CHalf is equivalent to Ancestor-(HhH)$_2$-Sec-CHalf (**Figure S1d**). HPLC chromatogram and mass spectrum demonstrating successful NCL of the half peptides to yield Symmetric-(HhH)$_2$-(A29U) ($m_{calc}$ = 6754.9 Da, $m_{obs}$ = 6753.4 Da). **d**. HPLC chromatogram and mass spectrum demonstrating successful deselenization of Symmetric-(HhH)$_2$-(A29U) to yield Symmetric-(HhH)$_2$ ($m_{calc}$ = 6675.92 Da, $m_{obs}$ = 6674.0 Da). **e**. Binding of synthesized Symmetric-(HhH)$_2$ to 101 bp dsDNA as measured by SPR (25ºC, 20 μL/min flow rate). As with Ancestor-(HhH)$_2$, the kinetics are biphasic, and 1000 s were at some concentrations insufficient to reach steady state.

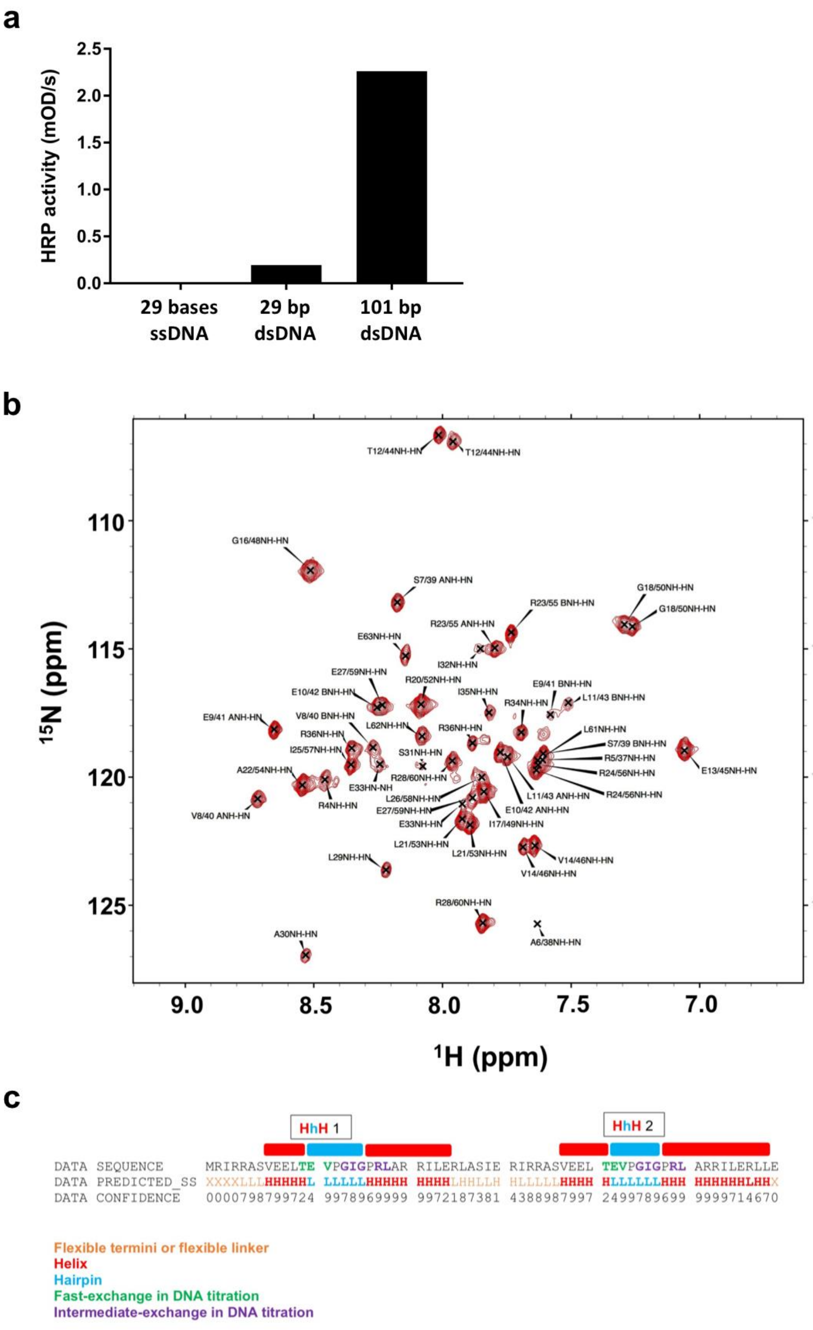


**Figure S3. Characterization of Primordial-(HhH)$_2$-Arg**. **a**. Binding of *E. coli* expressed, 6xHis-tagged Primordial-(HhH)$_2$-Arg to multiple dsDNA sequences as assayed by ELISA. **b**. NMR-HSQC spectrum reporting peak assignments of *E. coli* expressed, tag-free Primordial-(HhH)$_2$-Arg. **c**. TALOS+ neural network secondary structure prediction based on backbone chemical shift assignments. Secondary structure prediction is missing or low confidence for linker and N-terminal residues due to broadened spectra from flexible conformations.

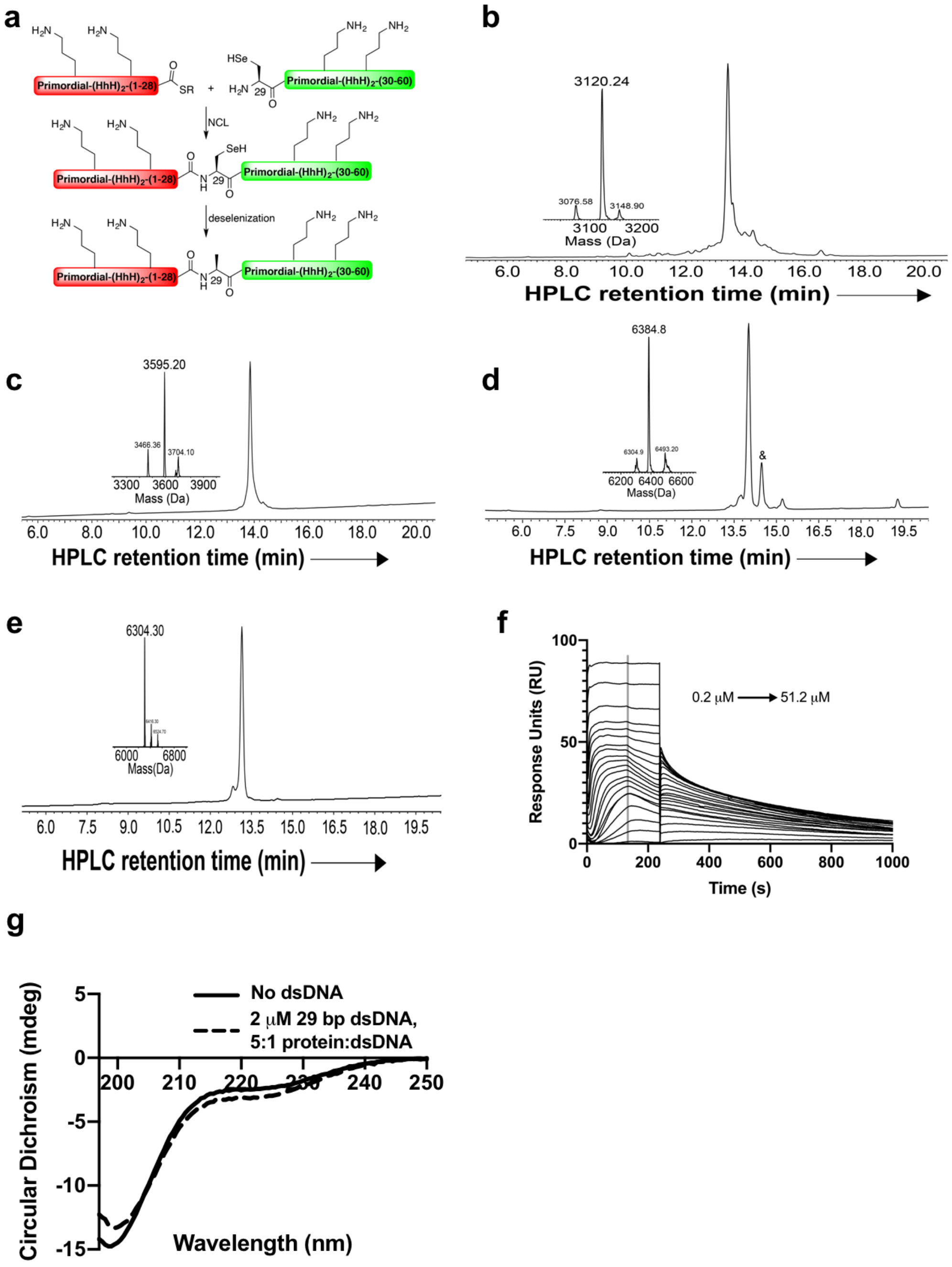


**Figure S4. Synthesis and characterization of Primordial-(HhH)$_2$-Orn**. **a**. Chemical protein synthesis scheme for Primordial-(HhH)$_2$-Orn. Because the protein was too long for a single solid phase synthesis reaction, two half-peptides were synthesized and then joined using an NCL and deselenization approach. In this case, the N-terminal half-peptide bears the C-terminal thioester surrogate Nbz moiety (NHalf-COSR) and the C-terminal peptide bears an N-terminal selenocysteine (Sec, U) residue (Sec-CHalf). After peptide ligation, the selenocysteine residue is deselenized to yield alanine. **b**. HPLC chromatogram and mass spectrum demonstrating successful synthesis of Primordial-(HhH)$_2$-Orn-NHalf-COSR ($m_{calc}$ = 3121.72 Da, $m_{obs}$ = 3120.24 Da). **c**. HPLC chromatogram and mass spectrum demonstrating successful synthesis of fragment Primordial-

$(HhH)_2$-Orn-Sec-CHalf ([M+Na] $m_{calc}$ = 3464.96 Da, $m_{obs}$ = 3466.36 Da; [M+TNP] $m_{calc}$ = 3596.11 Da, $m_{obs}$ = 3595.20 Da). **d**. HPLC chromatogram and mass spectrum demonstrating successful NCL of the half peptides to yield Primordial-$(HhH)_2$-Orn-(A29U) ($m_{calc}$ = 6386.53 Da, $m_{obs}$ = 6384.80 Da). Column impurities are denoted with an ampersand (&). **e**. HPLC chromatogram and mass spectrum demonstrating successful deselenization of Primordial-$(HhH)_2$-Orn-(A29U) to yield Primordial-$(HhH)_2$-Orn ($m_{calc}$ = 6307.57 Da, $m_{obs}$ = 6305.50 Da). **f**. Binding of synthesized Primordial-$(HhH)_2$-Orn to 101 bp dsDNA as measured by SPR (25 ºC, 20 μL/min flow rate, 250 s contact time). The grey line denotes the time points taken to generate the binding curves in **Figure 3c**. **g**. A small yet consistent increase in the CD signal that reports α-helicity is observed upon mixing Promoridal-$(HhH)_2$-Orn with 29 base pair dsDNA, suggesting folding upon binding. This interpretation is bolstered by the more notable increase in α-helicity observed for the statistically guanidated variants which also show more avid binding compared to Primordial-$(HhH)_2$-Orn (**Figure 3e**). Plotted spectra are background-subtracted to remove the signal associated with dsDNA.

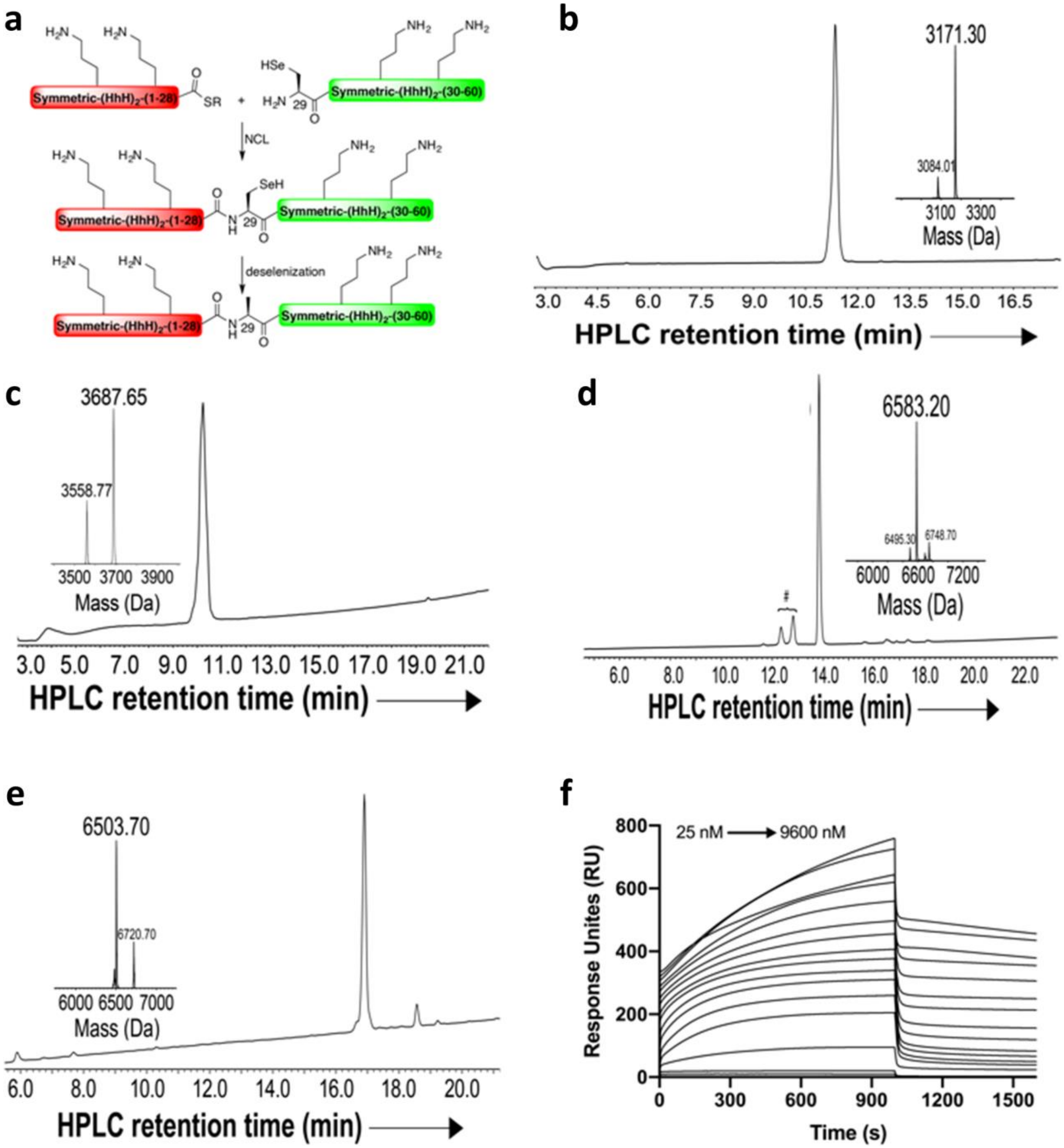


**Figure S5. Synthesis and characterization of Symmetric-(HhH)2-Orn**. **a**. Chemical protein synthesis scheme for Symmetric-(HhH)2-Orn. Because the protein was too long for a single solid phase synthesis reaction, two half-peptides were synthesized and then joined using an NCL and deselenization approach. The N-terminal half-peptide bears the C-terminal thioester surrogate Nbz moiety (NHalf-COSR), and the C-terminal peptide bears an N-terminal selenocysteine (Sec, U) residue (Sec-CHalf). After peptide ligation, the selenocysteine residue is deselenized to yield alanine. **b**. HPLC chromatogram and mass spectrum demonstrating successful synthesis of Symmetric-(HhH)2-Orn-NHalf-COSR ($m_{calc}$ = 3171.75 Da, $m_{obs}$ = 3171.30 Da). **c**. HPLC chromatogram and mass spectrum demonstrating successful synthesis of fragment Symmetric-

$(HhH)_2$-Orn-Sec-CHalf ([M+Na] $m_{calc}$ = 3557.96 Da, $m_{obs}$ = 3558.77 Da; [M+TNP] $m_{calc}$ = 3689.11 Da, $m_{obs}$ = 3687.65 Da). **d**. HPLC chromatogram and mass spectrum demonstrating successful NCL of the half peptides to yield Symmetric-$(HhH)_2$-Orn-(A29U), in which position 29 is selenocysteine ($m_{calc}$ = 6586.56 Da, $m_{obs}$ = 6583.20 Da). Column impurities are denoted with a hash (#). **e**. HPLC chromatogram and mass spectrum demonstrating successful deselenization of Symmetric-$(HhH)_2$-Orn-(A29U) to yield Symmetric-$(HhH)_2$-Orn ($m_{calc}$ = 6507.60 Da, $m_{obs}$ = 6503.70 Da). **f**. Binding of synthesized Symmetric-$(HhH)_2$-Orn to 101 bp dsDNA as measured by SPR (25 ºC, 20 μL/min flow rate, 1000 s contact time).

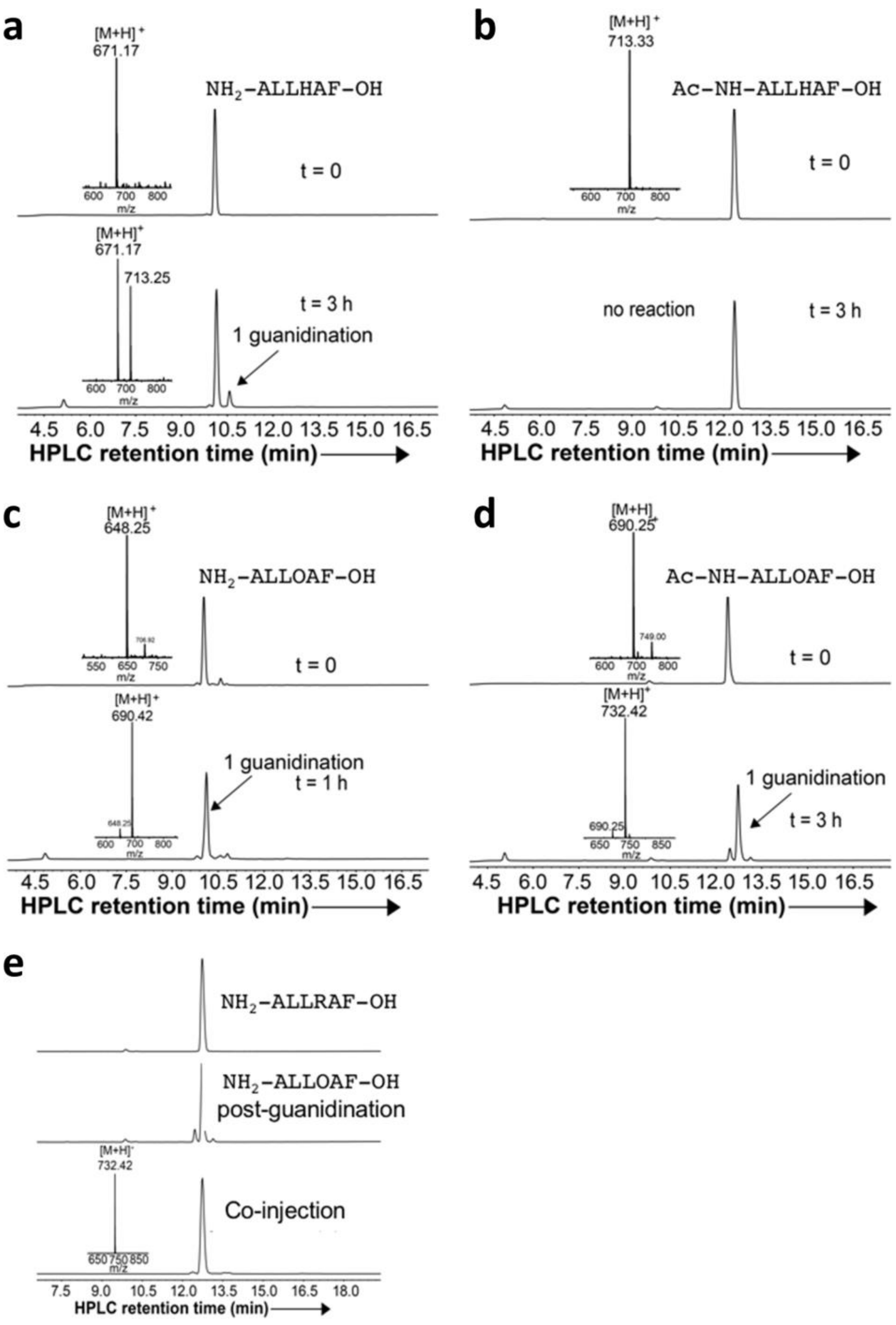


**Figure S6. Assessing the kinetics, efficiency, and specificity of the guanidination reaction**. **a**. To determine the reactivity of the N-terminus, and confirm that His residues are not modified by the guanidination reaction, $NH_2$-ALL**H**AF-OH was synthesized (upper HPLC chromatogram; $m_{calc}$ = 671.39 Da, $m_{obs}$ = 671.17 Da) and subjected to the guanidination reaction (lower HPLC chromatogram). Slow, single guanidination of the peptide was observed after three hours ($m_{calc}$ = 713.41 Da, $m_{obs}$ = 713.25 Da). **b**. No guanidination of Ac-NH-ALL**H**AF-OH was observed after 3 hours, as indicated by analytical HPLC and ESI-MS analysis ($m_{calc}$ = 713.40 Da, $m_{obs}$ = 713.33 Da), indicating that the slow, single guanidination of the peptide described in Panel A occurs at the amino-terminus. **c**. Synthesis of $NH_2$-ALL**O**AF-OH (upper HPLC chromatogram; $m_{calc}$ = 648.41 Da, $m_{obs}$ = 648.25 Da). Near complete guanidination was achieved within one hour, resulting in a single major peak ($m_{calc}$ = 690.43 Da, $m_{obs}$ = 690.42 Da). Preferential guanidination of ornithine relative to the N-terminus is likely due to steric effects[2–4]. **d**. Synthesis of Ac-NH-ALL**O**AF-OH (upper HPLC chromatogram; $m_{calc}$ = 690.42 Da, $m_{obs}$ = 690.25 Da). A single major peak corresponding to the conversion of ornithine to arginine was observed (lower HPLC chromatogram; $m_{calc}$ = 732.44 Da, $m_{obs}$ = 732.42 Da). **e**. The peptide $NH_2$-ALL**R**AF-OH (*i.e.*, the expected guanidination product of $NH_2$-ALL**O**AF-OH) was synthesized (upper HPLC chromatogram), and showed virtually identical retention time as the guanidination product of $NH_2$-ALL**O**AF-OH (middle HPLC chromatogram). When the two samples were mixed and co-injected, the resulting HPLC chromatogram contained a single major peak, the mass of which matched that of $NH_2$-ALL**R**AF-OH ($m_{calc}$ = 732.44 Da, $m_{obs}$ = 732.42 Da), indicative of efficient and selective conversion to arginine. Analytical HPLC was performed on an XSelect C18 column.

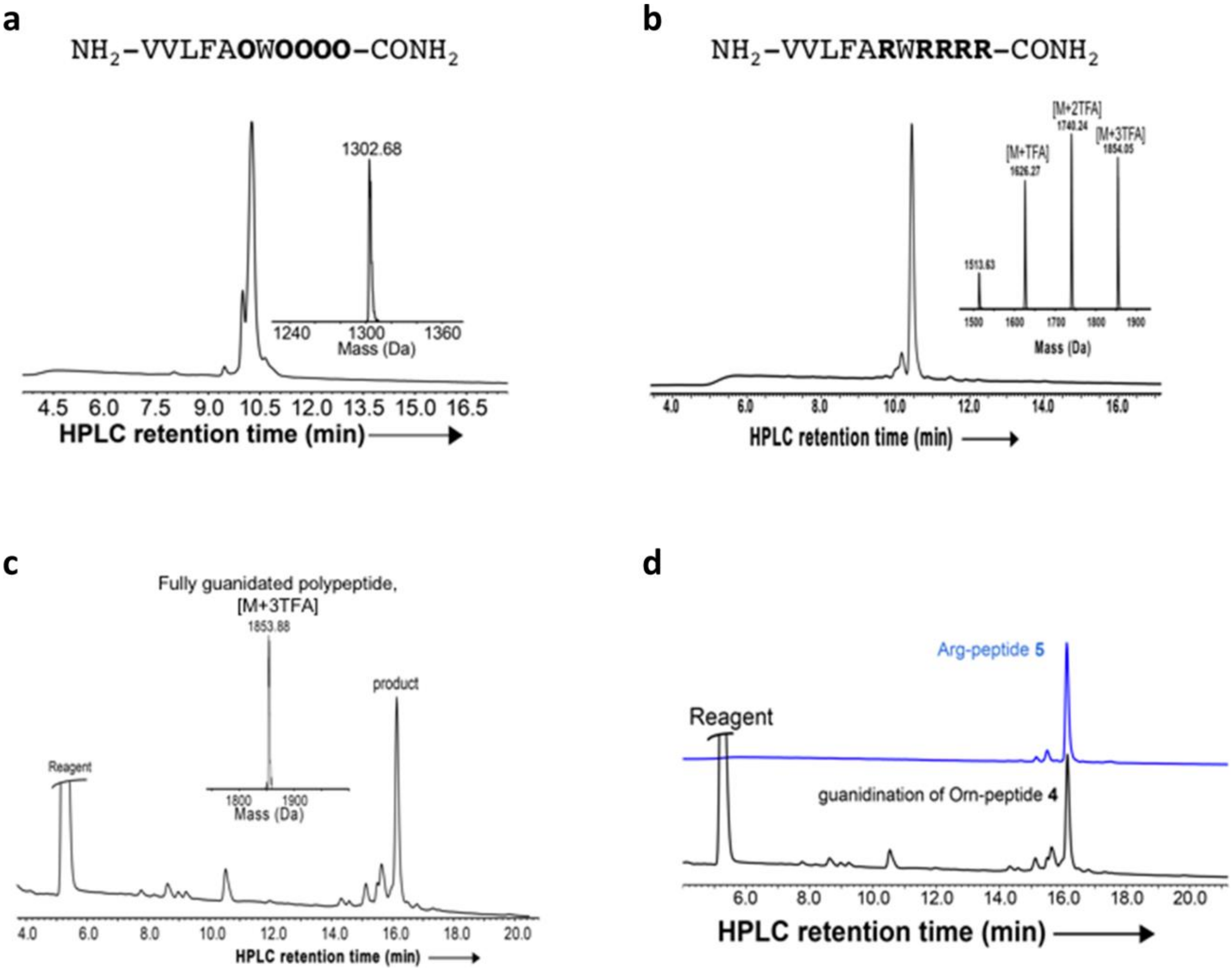


**Figure S7. Optimizing the guanidination of polypeptides with multiple ornithine residues**. **a**. To confirm that guanidination of adjacent ornithine residues can be achieved, NH$_2$-VVLFA**O**W**OOOO**-OH was synthesized ($m_{calc}$ = 1302.83 Da, $m_{obs}$ = 1302.68 Da). The small shoulder peak is the result of incomplete coupling between ornithine and tryptophan. **b**. The corresponding arginine containing peptide, NH$_2$-VVLFA**R**W**RRRR**-OH, was synthesized for comparison ($m_{calc}$ = 1513.87 Da, $m_{obs}$ = 1513.63, as well as TFA salts). **c**. After 2 hours of reaction, the ornithine peptide (NH$_2$-VVLFA**O**W**OOOO**-OH) showed complete guanidination, with all of its 5 ornithine residues converted to arginine ([M+3TFA] $m_{calc}$ = 1852.92 Da, $m_{obs}$ = 1853.88). **d**. The post-guanidination ornithine peptide and the synthetic arginine peptide have the same retention times on an HPLC XSelect C18 column.

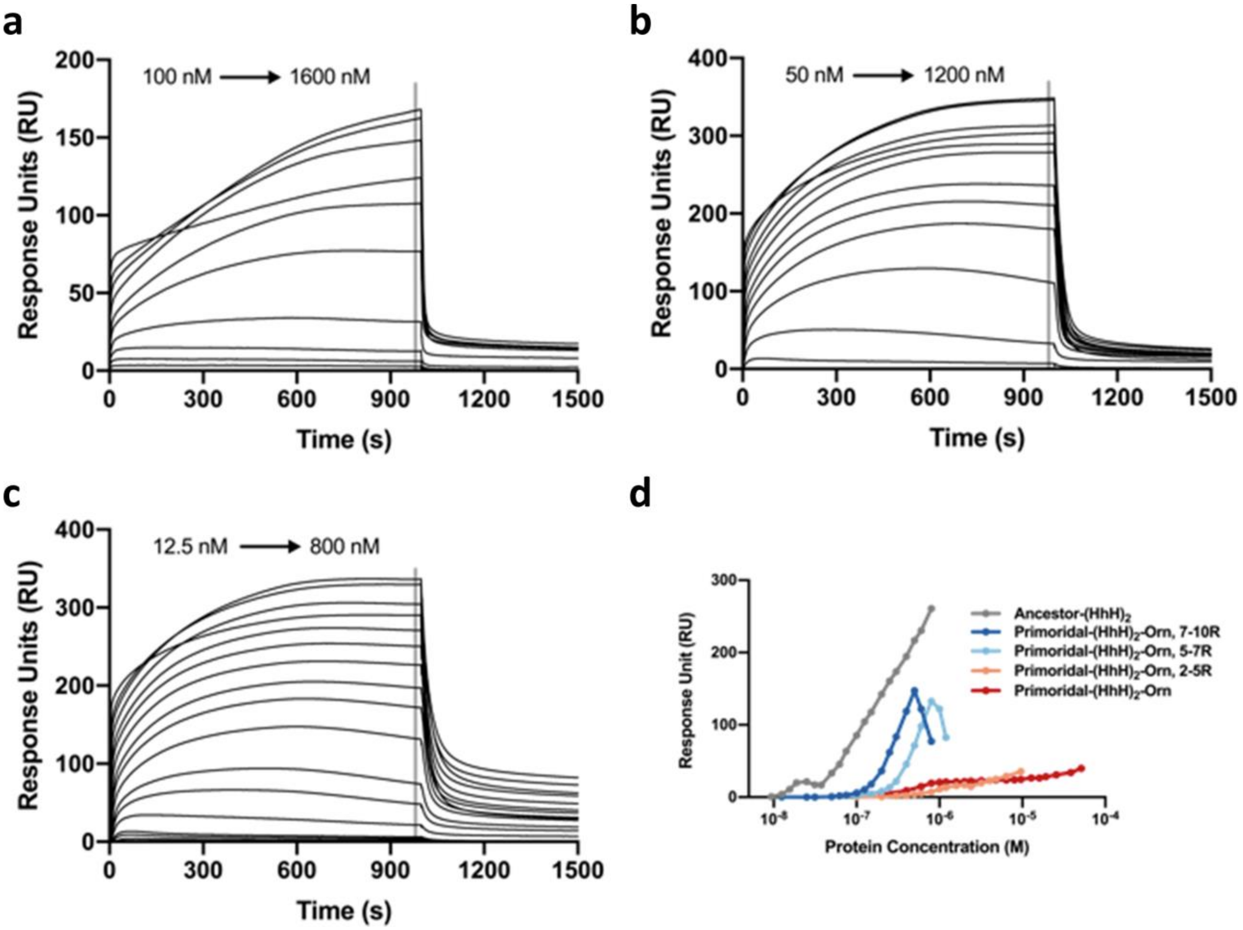


**Figure S8. Binding of guanidinated variants of Primordial-(HhH)$_2$ to dsDNA as measured by SPR.** **a**. Sensograms of binding of Primordial-(HhH)$_2$, 2-5R to 101 base pair dsDNA. **b**. Binding of Primordial-(HhH)$_2$, 5-7R to 101 base pair dsDNA. **c**. Binding of Primordial-(HhH)$_2$, 7-10R to 101 base pair dsDNA. As with other constructs, the association kinetics are biphasic, and in many runs (**Panels a-c**) steady state was not reached within 1000 s (see legend of **Figure 1**). The grey lines denote the time points taken to generate the steady-state binding curve in **Figure 3c**). **d**. Steady-state binding of guanidinated variants of Primordial-(HhH)$_2$ to 29 base pair dsDNA as measured by SPR (raw sensograms not shown). SPR experiments were performed at 25 ºC with a 20 μL/min flow rate and 1000 s contact time, except for Primordial-(HhH)$_2$-Orn, which required a contact time of 250 s.

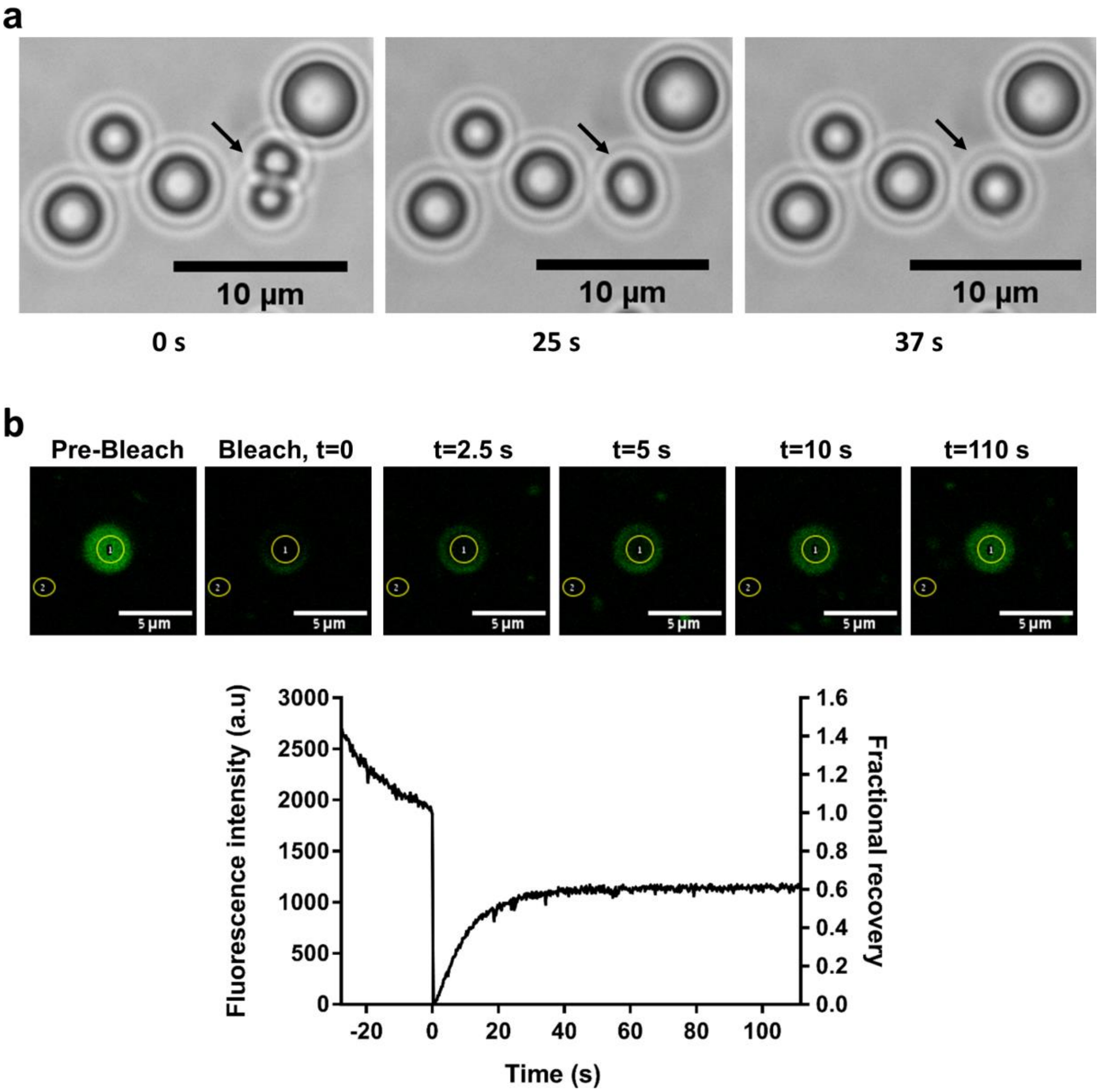


**Figure S9. Phase separation of Precursor-Arg with PolyU**. **a**. Droplets coalescence occurs over short time scales, confirming their fluid nature, as observed by transmission microscopy using a 100x oil immersion objective. Reaction conditions are the same as in **Figure 4** in the main text. **b**. FRAP analysis of fluorescent droplet using an inverted confocal microscope (Olympus IX81) with 60x oil immersion objective. Bleaching was performed inside the droplet, and recovery was followed over time. Fluorescence intensity was background normalized by subtracting the signal outside the droplet (area 2) from the signal inside the droplet (area 1). Fractional recovery was normalized by signal immediately prior to bleaching. The reaction mixture contains 1.0 mg/mL polyU and 208 μM Precursor-Arg (200 μM unlabeled protein plus 8 μM fluorescein-labeled protein).

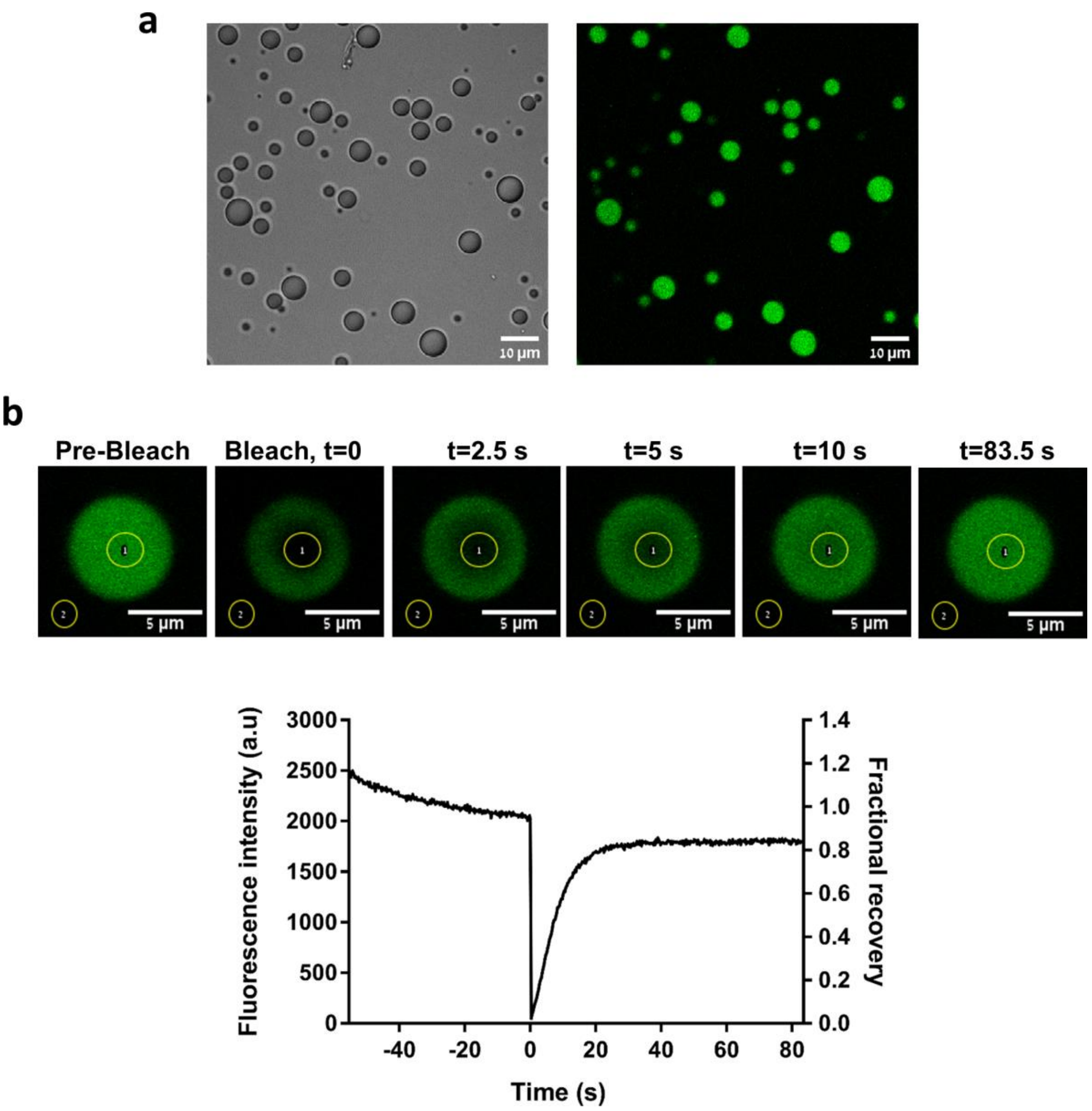


**Figure S10. Phase separation of Ancestor-(HhH)₂.** **a**. Ancestor-(HhH)$_2$ (200 µM unlabeled protein plus 4 µM fluorescein labeled protein) and polyU (1 mg/mL) form droplets upon mixing, as observed by transmission and fluorescence microscopy. **b**. FRAP analysis of fluorescent droplet on inverted confocal microscope with 60x oil immersion objective. Bleaching was performed inside the droplet and recovery was followed over time. Fluorescence intensity was background normalized by subtracting the signal outside the droplet (area 2) from the signal inside the droplet (area 1). Fractional recovery was normalized by the signal immediately prior to bleaching. Images were taken using inverted confocal microscope (Olympus IX81) with 60x oil immersion objective.

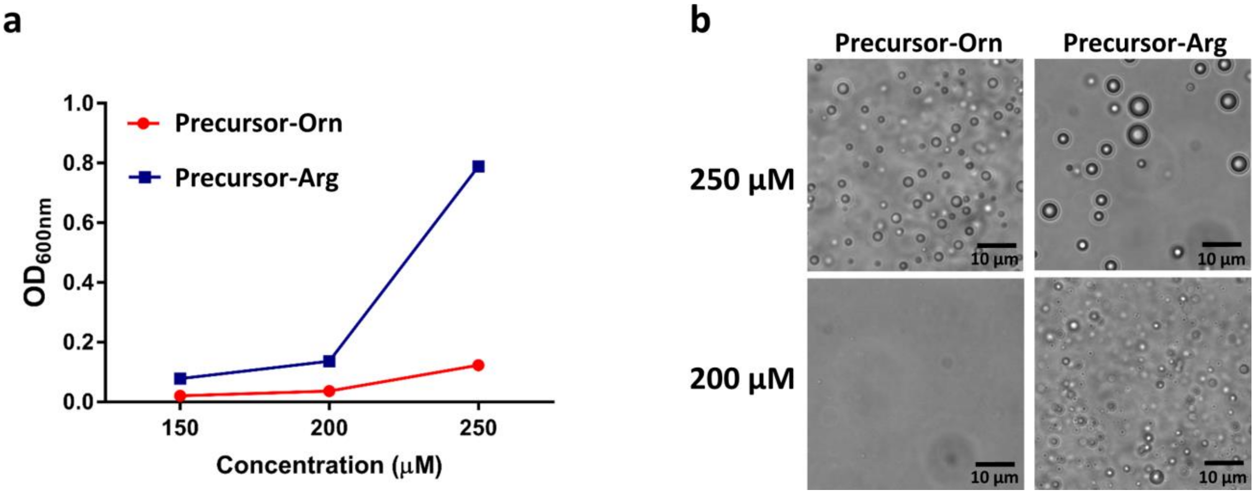


**Figure S11**. **Precursor-Arg forms larger droplets at lower concentrations than Precursor-Orn**. **a**. The dependence of phase separation on polypeptide concentration in the presence of 1 mg/mL PolyU. The extent of phase separation was assessed by turbidity measurements at 600 nm after 5 minutes of incubation at room temperature. **b**. Transmission micrographs of coacervates formed by 200 µM or 250 µM polypeptide and 1 mg/ml polyU.

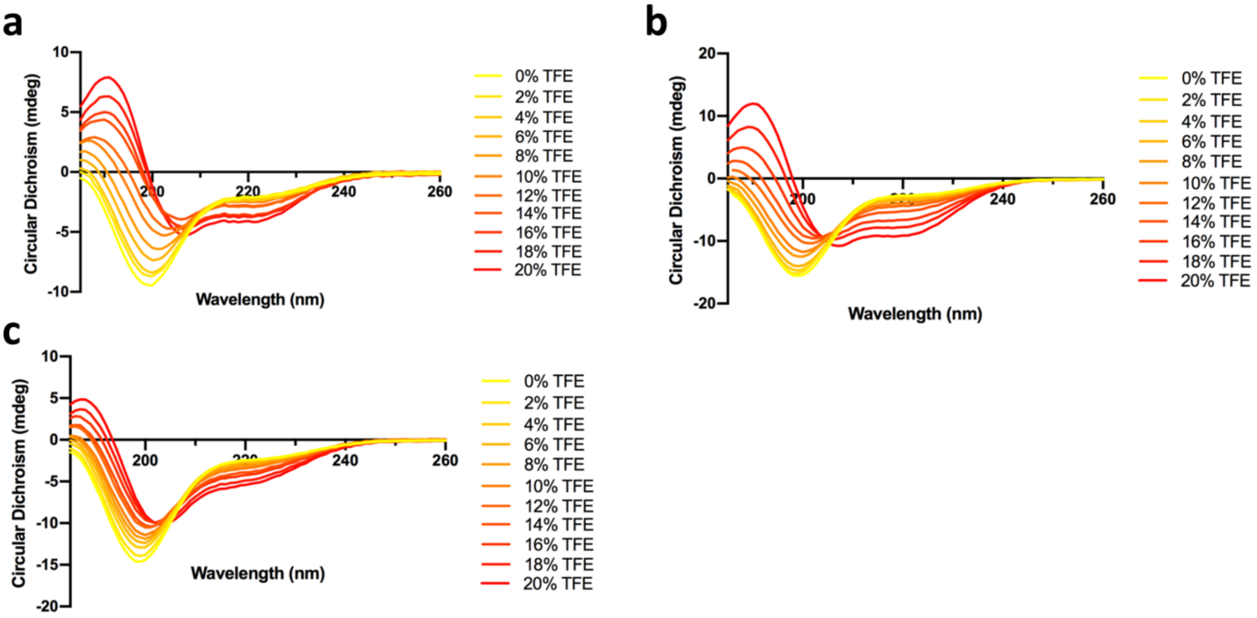


**Figure S12. Circular dichroism spectra from trifluoroethanol (TFE) titrations**. **a**. Precursor-Arg spectra. **b**. Scrambled 1 spectra. **c**. Scrambled 2 spectra. Each curve represents the average of two spectra after correction for dilution due to titration with TFE and subtraction of the buffer signal.

#### Materials and Methods

**Reconstruction of Ancestor-$(HhH)_2$.** Sequences for each of the $(HhH)_2$ protein families were collected from Pfam[5] and filtered to 50% identity using CD-HIT[6]. Initial alignments were generated with MAFFT-LINSI[7] then curated manually (sequence alignment is **Supplemental File 1**). An unrooted phylogentic tree for the $(HhH)_2$ protein family was built using IQ-TREE[8], with automatic model selection. The tree was found to be monophyletic with respect to the four known $(HhH)_2$ protein families (**Figure 1b**). Ancestral state inference was performed with CODE-ML from the PAML Suite package[9]. Plots of posterior probabilities were generated using R package GGLogo and are shown in **Figure 1c**.

**Protein expression in *E. coli* and purification**. Synthetic genes were ordered from Twist Bioscience (www.twistbioscience.com) and cloned into a pET21 vector, yielding variants with a C-terminal 6xHis tag. Mutant constructs were generated following Wang and Malcolm[10]. All constructs were verified by Sanger sequencing. Transformed BL21(DE3) cells were induced at $OD_{600}$ = ~0.6 with 1 mM IPTG. Induced cells were shake-incubated at 20 °C overnight. Cell pellets were collected by centrifugation and frozen at -20 °C for storage. The purification protocol was developed throughout this study as more variants were expressed and tested; the optimized protocol is as follows: Cell pellets from 250 mL cultures were resuspended in 30 mL of 100 mM NaCl, 50 mM Tris/HCl pH 7.5, 1:10,000 dilution of Benzonase, 1:10,000 dilution of rDNase1, 0.25x Protease Inhibitor Cocktail for Histidine-tagged proteins (Sigma-Aldrich), and 0.3 mg/mL and shake-incubated at 37 °C for 90 minutes. The lysates were then cooled on ice for 15 minutes and sonicated at 30 % power for 5 minutes, with a 30 s cool-down period after every 30 s of sonication. After cell lysis, the samples were spiked with 1 M NaCl and 5 mM imidazole and gently rocked for 15 min at room temperature, as high concentrations of NaCl promoted the solubility of some of the $(HhH)_2$ proteins. Pellets were clarified by centrifugation at 10,000 RCF for 1 hour and passed through a 0.45 µm sterile filter. Clarified lysate was applied to 3 mL packed NiNTA resin pre-equilibrated in 5 mM imidazole, 1 M NaCl, 50 mM Tris/HCl, pH 8.0 ("purification buffer"). After sample application, the resin was washed with 25 mL of purification buffer. Some $(HhH)_2$ constructs were copurified with dsDNA; to remove it, the resin was washed first with 25 mL of purification buffer containing 4M GuHCl, then with 50 mL of purification buffer containing 3 M NaCl. Nonspecifically bound proteins were removed by washing with purification buffer spiked with 25 mM imidazole. The bound $(HhH)_2$ protein was eluted by 5 mL of 850 mM imidazole, 50 mM Tris/HCl, 150 mM NaCl. Samples were then concentrated to ~0.5 mL and analyzed by SDS-PAGE gel to determine their purity and estimate the protein concentration. The absence of residual dsDNA was confirmed by a Qubit Fluorometer calibrated for dsDNA (ThermoFisher Scientific).

**Enzyme-linked immunosorbent assay (ELISA)**. Binding to DNA was measured by an ELISA assay. DNA sequences used for the binding assay can be found in **Table S5**. The arbitrarily chosen promoter region of the *nrdH* gene (Gene ID: 947161) was amplified from the *E. coli* genome, then subjected to a second round of PCR using a single 5'-biotinylated primer. The 101 base pair PCR product was purified with a QIAquick PCR purification Kit (Qiagen), eluted in water, and the DNA concentration was determined by measuring $A_{260nm}$ (NanoDrop 2000, ThermoScientific). Other dsDNA constructs were ordered as oligonucleotides (Integrated DNA Technologies) and annealed with their complementary oligonucleotide by heating to 95 °C, followed by slow cooling down to room temperature at a rate of 2 °C per min. Streptavidin-coated plates (StreptaWell, Roche) were incubated for 30 min with 100 µl solutions of either 1 µM biotin, 1.5 ng/µl 29 bases of biotinylated ssDNA, 1.5 ng/µl 29 base pair biotinylated dsDNA, or 1.5 ng/µl 101 base pair biotinylated dsDNA (**Table S5**) in binding buffer (50 mM Tris/HCl pH 8.0, 150 mM NaCl). The wells were washed extensively with binding buffer, then blocked with binding buffer spiked with 1 mg/mL BSA. Coated wells were incubated for 45 min with proteins in binding buffer plus 1 mg/mL BSA. The unbound proteins were washed, and 100 µl of 1 µg/ml HRP-labeled mouse anti-His antibody (200 µg/ml, Santa Cruz Biotechnology) was added. Following 45 min incubation, the wells were washed, the substrate 3,3',5,5'-tetramethybenzidine (TMB; ES001, Millipore), was added, and $OD_{650nm}$ was monitored. ELISA binding data are presented as the rate of increase in $OD_{650nm}$ with the background (wells coated with biotin) subtracted (null in nearly all cases).

**Materials for total chemical protein synthesis**. Buffers were prepared using MilliQ water (Millipore, Merck). Ultrapure guanidinium chloride (Gn·HCl, MP Biomedicals, LLC, France) was used in all ligation reactions. $Na_2HPO_4 \cdot 12H_2O$, tris(2-carboxyethyl)phosphine hydrochloride (TCEP·HCl), ethanedithiol (EDT), triisopropylsilane (TIPS), *D,L*-dithiothreitol (DTT), 2,2′-Dithiobis(5-nitropyridine) (DTNP), sodium ascorbate and 4-mercaptophenylacetic acid (MPAA) and methyl 3-mercaptopropionate (MMP) were purchased from Sigma-Aldrich (Rehovot, Israel). 1-guanyl-3,5-dimethyl pyrazole nitrate (the guanidination reagent) was purchased from Chem-Impex. All Fmoc-amino acids were obtained from CS Bio Co. (Menlo Park, CA) or Matrix Innovation (Quebec City, Canada), with the following side chain protecting groups: Arg(Pbf), Glu(OtBu), Gly(OtBu), Ser(tBu), Thr(tBu), Tyr(tBu), Lys(Boc), His(Trt), Trp(Boc). (Pbf = 2,2,4,6,7- pentamethyl-2,3-dihydrobenzofuran-5-sulfonyl). Nα-Fmoc-Nδ-Boc-L-Ornithine (shortly Fmoc-Orn(Boc)-OH, three letter code Orn, and one letter code O) was purchased from Chem-Impex. TentaGel® R RAM resin (loading 0.19 mmol/g), Fmoc-Phe-TentaGel® R PHB resin (loading 0.18 mmol/g) and Fmoc-Leu-TentaGel® were purchased from Rapp Polymer GmbH (Germany). 1-[Bis(dimethylamino)methylen]-5- chlorobenzotriazolium 3-oxide hexafluorophosphate, *N*,*N*,*N′*,*N′*-Tetramethyl-O-(6- chloro-1H-benzotriazol-1-yl)uronium hexafluorophosphate (HCTU) and Ethyl cyano(hydroxyimino)acetate (OxymaPure) were purchased from Luxembourg Biotechnologies Ltd. (Rehovot, Israel). All solvents: *N*,*N*-dimethylformamide (DMF), dichloromethane (DCM), acetonitrile (ACN), *N*,*N*-diisopropylethyl amine (DIEA), Trifluoroacetic acid (TFA) and piperidine (Pip) were purchased from Bio-Lab (Jerusalem, Israel) and were peptide synthesis, HPLC, or ULC-grade. Synthesis of Fmoc-Sec(Mob)-OH (selenocysteine three letter code is Sec, and one letter codes is U) was reported previously[11].

**High performance liquid chromatography (HPLC)**. Analytical reversed-phase (RP) HPLC analyses were performed on a Waters Alliance HPLC with UV detection (220 nm and 280 nm) using a XSelect C18 column (3.5 µm, 130 Å, 4.6 × 150 mm) or XBridge C4 column (3.5 µm, 4.6 × 150 mm). Preparative and semi-preparative RP-HPLC were performed on a Waters 150Q LC system using a XSelect C18 column (5 µm, 30 × 250 mm) or XBridge BEH300 C4 column (5 µm, 19 × 150 mm). Linear gradients of ACN (with 0.1 % TFA, buffer B) in water (with 0.1 % TFA, Buffer A) were used for all systems to elute bound peptides. The flow rates were 1 mL/min (analytical column heated at 30 °C), 10 mL/min (semi-preparative), and 20 mL/min (preparative).

**Electrospray ionization mass spectrometry (ESI-MS)**. ESI-MS was performed on LCQ Fleet Ion Trap mass spectrometer (Thermo Scientific). Peptide masses were calculated from the experimental mass to charge (m/z) ratios from the observed multiply charged species of each peptide. Deconvolution of the experimental MS data was performed with the help of MagTran v1.03 software.

**General procedure for Fmoc-solid-phase peptide synthesis (Fmoc-SPPS)**. Peptides were prepared with an automatic peptide synthesizer (CS136XT, CS Bio Inc. CA), except otherwise noted) typically on 0.25 mmol scales. Fmoc-amino acids (2 mmol in 5 mL DMF) were activated with HCTU (2 mmol in 5 mL DMF) and DIEA (4 mmol in 5 mL DMF) for 5 min, then allowed to couple for 25 min with constant shaking. Fmoc deprotection was carried out with 20 % piperidine in DMF (2 × 5 min). Selenocysteine was coupled using DIC/OxymaPure activation method[11]. After synthesis, the peptide-resins were washed with DMF, DCM and dried under vacuum. The dried peptide-resins were deprotected and cleaved using a TFA/water/thioanisole/ triisopropylsilane/ethanedithiol (92.5:1.5:1.5:1.5:1.5) cocktail for 4 h. The cleavage mixtures were filtered and TFA was evaporated with $N_2$-bubbling to a minimum volume, to which an eightfold volume of cold ether was added dropwise. The precipitated crude peptide was centrifuged (5000 rpm, 10 min), ether was removed, and the crude peptide was dissolved in ACN/water (1:1) containing 0.1 % TFA, further diluted to ca. 25 % ACN with water, and lyophilized.

**Preparation of C-terminal thioester peptides**. The C-terminal thioester peptides were synthesized first as Fmoc-Dbz-resin (0.25 mmol scale) on automated peptide synthesizer. Mono-

Fmoc-3,4-diaminobenzoic acid (Fmoc-Dbz-OH, 3 equiv)[1] activated with HCTU (3 equiv)/DIEA (6 equiv) in DMF was doubly coupled to the free amine of TentaGel® R RAM resin (0.19 mmol/g, 0.25 mmol scale) for 1 h, followed by double coupling of the first amino acid (2 x 1 h), while the N-terminal peptide was protected with Boc-protecting group prior to the Dbz to Nbz conversion[1]. After synthesis completion, the resin was washed with DCM and a solution of *p*-nitrophenyl chloroformate (5 equivalents) in DCM was added, shaken for 1 h at 25 °C and washed with DCM (3 × 5 mL) and DMF (3 × 5 mL) (repeated twice). Following this, the resin was washed with DCM and DMF and a solution of 0.5 M DIEA in DMF was added and shaken for additional 30 min to complete the cyclization and Nbz formation (repeated twice), and washed with DMF (3 × 5 mL). Peptide was deprotected and cleaved as described previously to give the crude peptide acylurea derivative, peptide-Nbz. The crude peptide-Nbz (100 mg, ~3 mM) was dissolved in phosphate buffer (100 mM, 6 M Gn·HCl, pH ~7) and treated with methyl 3-mercaptopropionate (MMP) (5% v/v) for 5-7 hours at room temp. The reaction was monitored by analytical HPLC (XSelect C18 column, 3.5 µm, 130 Å, 4.6 × 150 mm), and ESI-MS, and the peptide was purified by preparative HPLC.

**Native chemical ligation (NCL)**. The $(HhH)_2$ constructs were too long for a single SPPS reaction; thus, two half-peptides were synthesized then joined using native chemical ligation (NCL) and deselenization. In this case, the N-terminal half-peptide bears a C-terminal thioester (or thioester surrogate, such as *N*-acyl urea, Nbz[1]) moiety (NHalf-COSR) and the C-terminal peptide bears an N-terminal selenocysteine (Sec, U) residue (Sec-CHalf). Briefly, the NHalf-COSR peptide was dissolved in argon degassed PB buffer (100 mM $NaH_2PO_4$, 6 M Gn·HCl, 0.2 M MPAA, 0.05 M TCEP and 0.3 M sodium ascorbate, pH 7.3) and added to the Sec-CHalf peptide, yielding a final peptide concentration of about 1 mM. The progress of the reaction was followed by analytical HPLC (XBridge C4 column, 3.5 µm, 4.6 × 150 mm) with a gradient of 5-70 % buffer B over 20 min. The ligation was quenched after 4 h. The ligation product was purified by semi-preparative HPLC (XBridge BEH300 C4 column, 5 µm, 19 × 150 mm, method 25-50 % B over 45 min). The resulting peptide still contained a selenocysteine residue at the site of ligation.

**Deselenization reaction**. Sec-containing peptides were dissolved in 1 mL of argon degassed PB buffer (100 mM, 6 M Gn·HCl, 100 equivalents DTT, pH 7.3), and left for 30 min, upon which 100 equivalents of TCEP in 200 µL of the same argon degassed buffer were added[12,13]. The progress of the reaction was followed by analytical HPLC (XBridge C4 column, 3.5 µm, 4.6 × 150 mm) with gradient 5-70 % B over 20 min, and TCEP was added as needed to generate the desired deselenized product.

**Guanidination of ornithine-containing polypeptides**. Peptides (0.238 µmol) were dissolved in TDW to a concentration of about 2 mg/mL. Stock solution containing 50 mg of 1-guanyl-3,5 dimethyl pyrazole nitrate dissolved in 300 µL TDW was prepared (0.83 M). For guanidination reaction, ~6 equivalents per each ornithine residue present in the peptide were added to the 1 mL peptide solution. The pH was adjusted to 9.3 using 1M NaOH, and the reaction was incubated at 37 °C[14]. The progress of the reaction was followed by HPLC and ESI-MS, and 10 µL of reaction aliquots quenched with 20 µL of 0.1 % TFA in TDW were injected (XBridge C4 column, 3.5 µm, 4.6 × 150 mm) and eluted using a gradient of 5 % B in eluent A for 2 min then 5-50 % B over 20 min.

**Surface plasmon resonance (SPR)**. Binding to 29 base pair or 101 base pair dsDNA was monitored by surface plasmon resonance on a Biacore T200 system. Since the $(HhH)_2$ variants are positively charged at neutral pH, a C1 chip (GE Life Sciences), which carries less charge than the standard CM5 chips, was used. Streptavidin was conjugated to the chip surface using EDC/NHS chemistry, as directed in the C1 sensor chip manual. Approximately 2000 RU of streptavidin was stably conjugated to the chip surface, which was then blocked by ethanolamine. Subsequently, 300 RU of biotinylated 29 base pair or 101 base pair dsDNA was stably associated to the surface. Prior to data collection, a normalization cycle followed by three priming cycles were run to stabilize the instrument. Binding assays were performed in 50 mM Tris, 150 mM NaCl, 0.02 % Tween-20, pH 7.5 ("SPR binding buffer") using a flow rate of 20 μL/min at 25 °C. In most cases, 1000 s contact times were required to achieve steady-state binding, the notable exception being Primordial-$(HhH)_2$-Orn, which required only 250 s of contact time. Regeneration of the chip surface was achieved by a 60 s injection of 2 M NaCl in SPR Binding Buffer.

**Circular dichroism**. CD spectra were collected on a Chirascan circular dichroism spectrometer (Applied Photophysics). Samples containing 10 μM protein in 5 mM Tris/HCl, 25 mM NaCl, pH 7.5 were loaded into a 1 mm pathlength quartz cuvette. Spectra were collected from 190-260 nm with a data pitch of either 0.5 or 1 nm at 25 °C and a slit width of 1 nm. The PMT voltage during measurement was kept below 700 V, and data points exceeding this value were discarded. All reported spectra are buffer subtracted. In the case of dsDNA titration experiments, the spectrum of the dsDNA alone was subtracted from that of the sample containing both dsDNA and protein. Because dsDNA absorbs in the far UV, concentrations of both the dsDNA and the protein were kept relatively low (2 μM and 10 μM, respectively) and samples with a large excess of dsDNA could not be studied. Reliable data could only be collected down to about 195 nm, a region where both the protein and the dsDNA exhibit significant circular dichroism, though of opposite sign. At these low wavelengths, the circular dichroism signal of the protein was about two-fold greater than that of the dsDNA.

**NMR experiments**. All NMR experiments were performed at 25 °C on Bruker Avance 800 MHz spectrometer equipped with TXI triple resonance cryogenic probes. All NMR data were processed using Topspin (www.bruker.com) and analyzed using Sparky[15]. All NMR samples were prepared to 200 μM in 95 % $H_2O$/5 % $D_2O$ 20 mM Tris-D11 (Cambridge Isotope Laboratories) at pH 7.5 and 250 mM NaCl. High salt concentration and relatively low protein concentration were required to minimize sample precipitation. For backbone resonance assignments of unbound Primordial-$(HhH)_2$-Arg, 2D $^{15}N$/$^{13}C$ HSQCs, and 3D HNCO, HNCA, HNCACB, CBCA(CO)NH, HN(CO)CA, HN(CO)CACB were collected. Despite duplication of the HhH motif, nearly all HN-NH correlations were obtained, except for terminal residues that are flexible and the linker regions, where residues were broadened by exchange. Secondary structure was predicted by uploading N, HN, C, CA, CB, and HA chemical shifts TALOS+[16].

$^{1}H$-$^{15}N$ HSQC titrations were collected in tandem by first optimizing conditions with unbound protein, followed by molar ratio titrations of a 12 base pair dsDNA fagment (**Table S5**). DNA base-pairing was confirmed by detection of imino resonance in $H_2O$ NOESY spectra.

**$^{13}C$, $^{15}N$ protein preparation**. An insert encoding the solubility tag GB1[17] linked by TEV tag to Primordial-$(HhH)_2$-Arg was cloned into pET-21a, an *E. coli* plasmid expression vector (Merck Millipore) with a C-terminal 6xHis tag and TEV protease cleavage site. Plasmids were then transformed into chemically competent *E. coli* Rosetta cells (New England Biolabs) grown in M9 medium supplemented with 1 g/L $^{15}NH_4Cl$ and 2 g/L $^{13}C$-glucose. Protein expression was then induced with 1 mM IPTG at 20 °C. After overnight expression, cells were collected, lysed, and the protein purified by Ni-affinity which included a 4 M guanidium hydrochloride wash to ensure removal of non-specifically bound DNA. The GB1-tag was removed by TEV protease cleavage (New England Biolabs) and separated by an additional Ni-affinity purification step. Sample aggregation was assessed by size-exclusion fast protein liquid chromatography (Superdex 75 10/300 GL, GE Healthcare). Exchange into Tris-D11 buffer was achieved by successive rounds of concentration and dilution using 10 kDa / 3 kDa MW cutoff Amicon® Ultra-15 Centrifugal Filter Units (Millapore-Sigma).

**Fluorescein labeling of peptides**. Equimolar amounts of peptide and NHS-fluorescein (ThermoScientific, 0.3-3 mM) were resuspended in 100 µL of acetonitrile (Bio-Lab ltd, Israel). The pH was adjusted to ~9 by addition of triethanolamine, and the reactions were incubated for 1 hour. Since the peptides were not soluble in acetonitrile, the unreacted NHS-fluorescein could be removed by washing the pellet several times in fresh acetonitrile until the solution was clear. After washing, the residual acetonitrile was removed by vacuum evaporation for 30 min (Concentrator plus, Eppendorf). Dried peptides were dissolved in water and the concentration of the fluorescein-labeled peptide was estimated by measuring $A_{493nm}$ (NanoDrop 2000, ThermoScientific).

**Phase separation**. Peptides and polyuridylic acid (polyU, Sigma, P9528) were dissolved in MilliQ water. Peptide concentrations were measured using the Pierce™ BCAProtein Assay Kit

(ThermoFisher Scientific), and a polyU stock solution was prepared at 10 mg/ml. Phase separation was induced by mixing: MES buffer, from a 10x stock, to a final of 50 mM MES pH 5.6, polyU to a final of 1.0-1.4 mg/ml, and the peptide to a final concentration of 190-240 µM. For imaging by fluorescent microscopy, fluorescein-labeled peptide was added (1-20 µM final concentration, in accordance with the degree of labelling). Microscopy glass slides were PEG-silanized as previously described[18]. Capillary channel slides for microscopy were custom-made using PEGylated slides (24 × 40 mm, thickness 0.13-0.16 mm), microscope slides (24 mm × 60 mm, thickness 0.15-0.19) and cover slips (24 x 24 mm, thickness 0.13-0.16 mm). Typically, 20-30 µl of phase separation reaction mixture was loaded into chambers and observed using an inverted microscope (Nikon Eclipse Ti-S, Japan) with 20x objective (Plan Fluor, 20x/0.45, Nikon, Japan) or with an oil immersed 100x objective (Plan Apo, 100x/1.40 oil, Nikon, Japan). Fluorescent images were taken with a GFP filter (λexc = 470 ± 20 nm, λem = 525 ± 25 nm) and analyzed using the Fiji platform[19].

**Fluorescence recovery after photobleaching (FRAP)**. Fluorescence recovery after photobleaching was measured for Ancestor-$(HhH)_2$ and Primordial-$(HhH)_2$-Arg coacervates, prepared as described above, and loaded into capillary slides. Coacervates were imaged using an Olympus IX81 inverted confocal microscope equipped with a 60x oil immersion objective (UPlanSApo 60x/1.35 Oil, Olympus). Photobleaching was achieved by the Tornado method, in which samples were excited with a 488 nm laser diode (excitation wavelength 488 nm, emission wavelength 520-550 nm) at 100 % power for 200 ms. The imaging time was 278.1 ms/frame. The fluorescence intensity was monitored as a function of time for both the bleached area and the background, using the Fiji platform, and the recovery of the bleached region was normalized against the background. Data were analyzed using Prism (GraphPad Software Inc.).
